# EM-Claw: An Experience-Grounded Language-Model Agent for Auto Cryo-EM Single-Particle Analysis

**DOI:** 10.64898/2026.11.23.768522

**Authors:** Hong Shan, Jun-rui Li, Zhong-liang Xing, Qing-ning Yuan, Kai Wang, Shi-qi Xu, Heng Liu, Li-hua Zhao, H. Eric Xu

## Abstract

Single-particle cryo-electron microscopy (cryo-EM) processing remains strongly dependent on expert judgement because difficult specimens often require dataset-specific strategies that cannot be captured by a fixed pipeline. Here we present EM-Claw, an experience-grounded multi-agent system for adaptive cryo-EM single-particle analysis. Rather than following a predefined workflow, EM-Claw retrieves processing precedents from previously solved datasets, consults external guidance, proposes and independently evaluates alternative strategies, executes CryoSPARC or RELION jobs, and revises the processing route according to quantitative and visual evidence. Human expertise can be introduced at any stage through natural-language instructions, while successful decisions and processing trajectories are retained as reusable experience. We evaluated EM-Claw on both publicly available cryo-EM datasets and experimental datasets collected in our laboratory, spanning soluble proteins, membrane proteins, GPCR complexes and small transporter monomers. Across these cases, EM-Claw adapted its processing strategies to specimen-specific challenges, including compositional heterogeneity, complex flexibility, preferred orientation and weak particle features. It produced reconstructions at higher resolution than previously published results (0.1∼1.3 Å improvement) or solved structures that are now being prepared for publication, with no or little human intervention, thereby substantially reducing the time, effort and field experience required for human decision-making.

## Introduction

Cryo-electron microscopy (cryo-EM) single particle analysis (SPA) now routinely yields near-atomic-resolution structures of macromolecules, and data collection has become largely automated. Converting the data into a structure, however, still depends on an expert making a long series of judgements: how to organize the pipeline, how to set the parameters of each job, which two-dimensional (2D) and three-dimensional (3D) classes to retain, whether the particle has symmetry, and how to analyse the resulting reconstructions. This dependence was recognized early: processing “frequently becomes a severe bottleneck, requiring expert intervention, prior structural knowledge, and weeks of calculations”^[1]^. In recent years, many tools have targeted specific steps in SPA, making automated SPA increasingly feasible. Examples include tools for micrograph curation and denoising^[2, 3]^, particle picking^[4–6]^, 2D-class ranking^[7, 8]^, particle-quality scoring^[9, 10]^, orientation diagnostics^[11, 12]^, correction of preferred orientation^[13, 14]^, and heterogeneity analysis^[15–17]^.

End-to-end systems that assemble multiple steps into a complete SPA workflow have also emerged, including RELION Schemes^[7]^, TranSPHIRE^[18]^, a Scipion-based unattended pipeline deployed at the ESRF^[19]^, CryoWizard^[10]^, and CryoSPARC repeat-target workflows^[20]^. These systems specify the overall route and parameter settings in advance and report failures rather than replanning. They therefore perform well on the types of data for which they were tuned, but they have no mechanism for finding a different route, or for improving themselves, when a dataset falls outside that range.

In practice, datasets differ in signal-to-noise ratio, junk content,sample purity, symmetry, conformational heterogeneity, and orientation distribution, and the appropriate processing route differs accordingly. An experienced practitioner does not simply execute a fixed recipe. Instead, the practitioner examines the micrographs, class averages, and maps; recognizes the situation; recalls how a similar dataset was solved; consults manuals and the literature; tests and figures out the solution; summarizes the experience for the next comparable case. By now, the automation for SPA has not yet captured this combination of perception, recall, adaptive experimentation, and accumulated experience.

Large language model (LLM) agents offer a potential route toward this form of flexible automation by approximating key aspects of expert reasoning within an iterative cycle of observation, interpretation, action, and evaluation. Coding agents such as Claude Code^[21]^ and OpenAI Codex^[22]^ illustrate this approach: they inspect code and execution results, reason about the next steps, make changes using tools, and revise their strategies based on validation feedback. Interleaving reasoning with tool use^[23, 24]^ has enabled agents to plan and, in some cases, execute experiments in chemistry^[25, 26]^, gene editing^[27]^, and biomedical research^[28–31]^. LLM agents have also been evaluated for operating scientific instruments, such as atomic force microscopes^[32]^. For cryo-EM processing, cryoAgent demonstrates the potential of LLM-based reasoning through a progressive architecture that combines reliable baseline processing, agent-guided optimization and heterogeneity analysis, and hypothesis-driven exploration^[33]^. A further direction is to give LLM agents a more central role in determining cryo-EM processing strategies, drawing on their ability to integrate diverse evidence, reason about alternative approaches, and adapt plans as new results emerge.

Here, we present EM-Claw, a multi-agent system in which LLM agents devise cryo-EM processing strategies in a manner analogous to human experts: examining the data, drawing on experience, composing and adapting a processing route accordingly, while accumulating reusable experience (Figure 1). EM-Claw is implemented in Claude Code and controls CryoSPARC^[1]^ and RELION^[34, 35]^ for data processing. It can operate fully unattended or be steered conversationally at any point during a run, and users can add external tools as needed. Using cryo-EM processing as a practical testbed, this work explores how LLM agents can be adapted to specialized scientific workflows that require domain knowledge, flexible decision-making and iterative evaluation.

**Figure 1.**
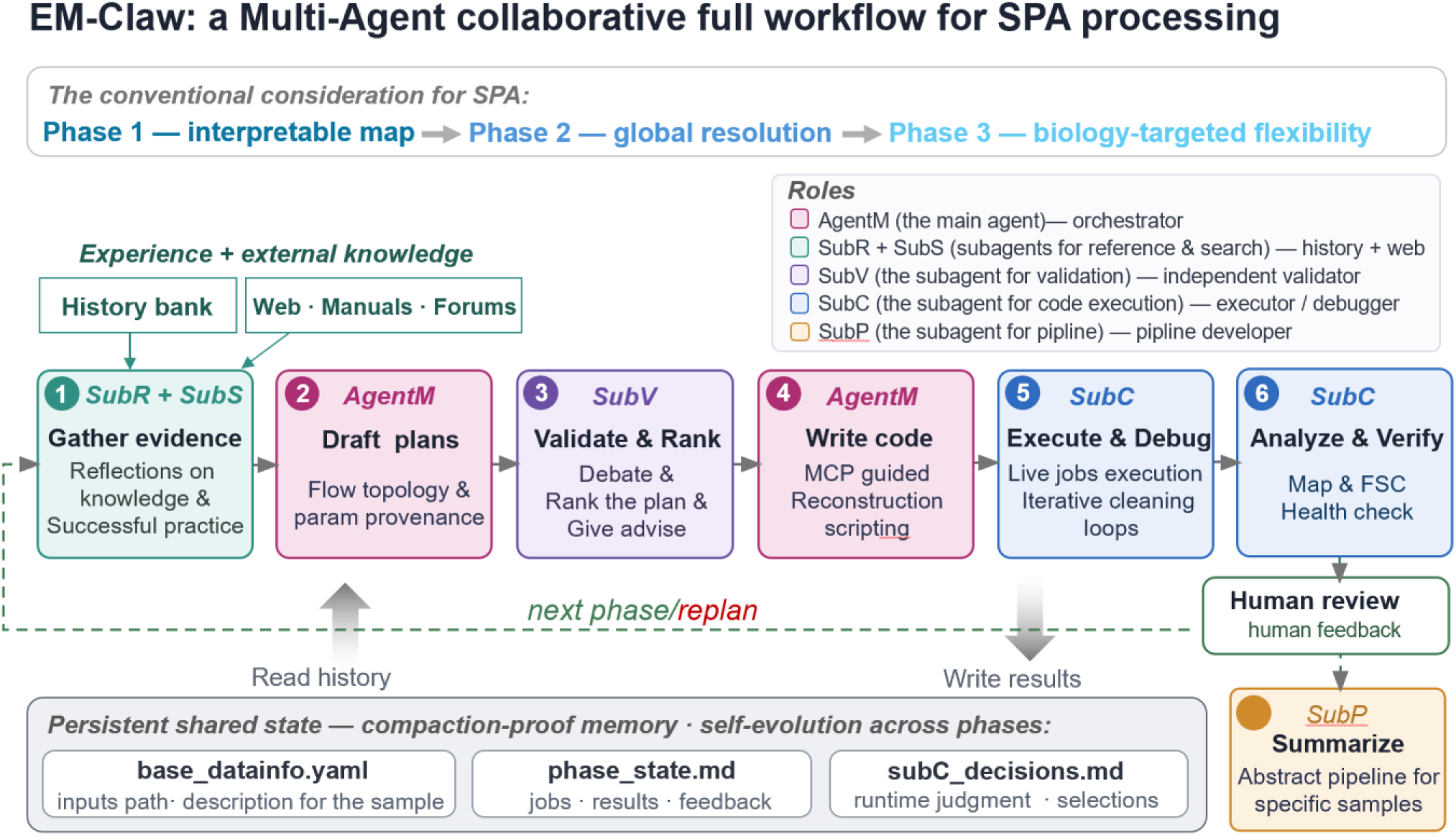
Multi-agent architecture of EM-Claw. An orchestrator (agentM) gathers precedents and external guidance through subR and subS, drafts alternative plans, obtains independent review from subV, translates the selected plan into executable code, and delegates execution and diagnosis to subC. Human review closes each phase and can trigger replanning. Dataset metadata, phase history, and run-time decisions are maintained in persistent files. subP distils completed projects into reusable pipelines outside the reconstruction loop.

## Materials and methods

### 1. System architecture

#### 1.1 Overview

EM-Claw is a multi-agent system that processes a cryo-EM single-particle dataset from raw movies or micrographs to a validated reconstruction (Figure 1). It is organized around a single principle: the processing strategy is composed for each dataset rather than selected from a predefined workflow. An orchestrator agent, agentM (the main agent), makes every strategic decision in accordance with the scientific objectives and constraints specified by the user. It delegates four recurring tasks to specialized agents, each invoked with an empty context and an explicit brief: subR (the subagent for reference works) retrieves precedents from previously completed processing projects; subS (the subagent for searching) searches the literature and software documentation for external guidance; subV (the subagent for validation) independently validates and ranks candidate plans; and subC (the subagent for code execution) executes and debugs jobs in a notebook-like manner. A fifth specialized agent, subP (the subagent for pipeline construction), is invoked outside the reconstruction loop to distil a completed project into a reusable pipeline that can be applied either within or independently of EM-Claw. Only the orchestrator can spawn agents; the specialized agents never spawn one another. This arrangement leaves a single agent accountable for the plan and makes every delegation auditable through its written brief. Compared with a single-agent design, this architecture isolates task-specific context and separates plan generation from independent review, reducing the orchestrator’s context burden during long processing runs while preserving coherent strategic control.

Processing advances in phases, each of which ends, by default, with a reconstruction for human review. A phase comprises one or more complete cycles: gathering evidence, drafting alternative plans, obtaining an independent ranking, making a decision, expressing the plan as code, executing it, and verifying the result. The human response to the delivered map then defines the problem for the next cycle. By convention, phase 1 seeks the first interpretable map, phase 2 seeks to improve global resolution, and phase 3 and subsequent phases focus on the density needed to answer the biological question, evaluated in the target region rather than solely by the global Fourier shell correlation (FSC).

Because a phase can span hours of computation and hundreds of tool calls, whereas each specialized-agent invocation begins with an empty context, persistent state is stored in files rather than in an agent’s conversational memory. Three file-based records carry this state (Figure 1, bottom). A project description records the dataset metadata, including data paths, acquisition optics, particle size and characteristics, symmetry, and paths to external tools. A phase record contains the human-facing history, with one entry per phase describing the jobs run, the result obtained, the human feedback, and a ledger that identifies branches already known to be unproductive. A decision log records the run-time selections made by the executing agent. In addition, a loop record contains one machine-readable row per round of every iterative stage, allowing an interrupted loop to be resumed and recovered.

#### 1.2 Evidence-grounded planning

Each phase begins with subR retrieving relevant precedent experience. During the initialization of the agent, for data safety and permission control, the precedents are first prepared outside the agent by deterministic scripts that read CryoSPARC project files or the CryoSPARC database. The agent then condenses them into a precedent bank for rapid consultation. Each digest retains the parameters that were explicitly set, the wiring of the job inputs, the particle counts, and, along the lineage that produced the project’s best map, the resolution gained at each step.

Two practices make retrieved precedents useful as evidence. First, they are matched by particle size and features, signal-to-noise ratio, and junk content. The last two are estimated from early 2D classification results and the fraction of picked particles retained for 3D analysis. Second, each summary records both parameter values and how processing steps are connected. This allows a plan to reproduce how multiple picking or selection branches were combined and when duplicates were removed.

A second agent, subS, retrieves external guidance. Its permitted sources depend on the phase. During the opening phase, it may cite only a curated set of authoritative references, ensuring that no uncommon setting enters the initial plan without provenance. During later phases, when the system is addressing a specific processing problem, the search is unrestricted. The agent is explicitly permitted to return no suggestion, reducing the incentive to manufacture a recommendation that its sources do not support.

The orchestrator then drafts three distinct plans by default. It constructs the plans by considering multiple precedents together: for example, one plan may adopt the particle-flow topology of one completed project and the parameter settings of another. It also emphasizes strategies should be tailored to the current dataset. This design provides alternative routes for comparison before committing computational resources.

#### 1.3 Independent evaluation of plans

The candidate plans are ranked by subV, a validator agent that uses a different model family from the planner and is accessed through a separate command-line interface. The validator checks each plan technically, so that a nonexistent job type, a mistyped parameter, or an incompatible connection can be detected before submission. It also checks each plan scientifically against the software manuals and the properties of the dataset. It returns a ranked list with a rationale for each plan and flags individual controversial parameters. The validator advises, but the orchestrator decides, because only the orchestrator holds the phase objective, the state of the run, and the user instructions.

Separating the agent that generates a plan from the agent that evaluates it is a deliberate control rather than an organizational convenience. Language models favour their own generations when acting as evaluators^[36]^ and do not reliably self-correct without external feedback^[37]^. The independent review helps identify technical errors and scientifically unsuitable choices before execution, while the use of a different model family may reduce shared blind spots between planning and validation. Explicit rationales and parameter flags make the assessment transparent and orchestrator’s control is preserved over the final decision.

#### 1.4 Execution and long-running jobs

The selected plan is expressed as code against the CryoSPARC application programming interface (API) or the RELION command line and is executed by subC. The executor works incrementally, in the manner of a notebook: it queues a job, inspects the result, and repairs failures at the smallest possible scope. If a plan cannot be implemented as written, it returns a repair request rather than improvising a new strategy so that e orchestrator retains control over the processing strategy.

Some jobs run for hours, and the executing agent does not receive a native notification when they finish. An agent that waits would therefore remain idle until its turn ended. Instead, the executor records the job identifier, the reason for queuing the job, and the intended next step, and then yields control. The orchestrator starts a background watcher that reports every terminal job state, including failure, and resumes the same executor when the job finishes.

#### 1.5 Autonomous judgement layer

To address the limitations of LLMs in interpreting three-dimensional structures and their two-dimensional projections, we developed approaches that combine computational scripts with vision models, enabling LLM agents to inspect and assess both 2D and 3D cryo-EM processing results (Figure 2).

**Figure 2.**
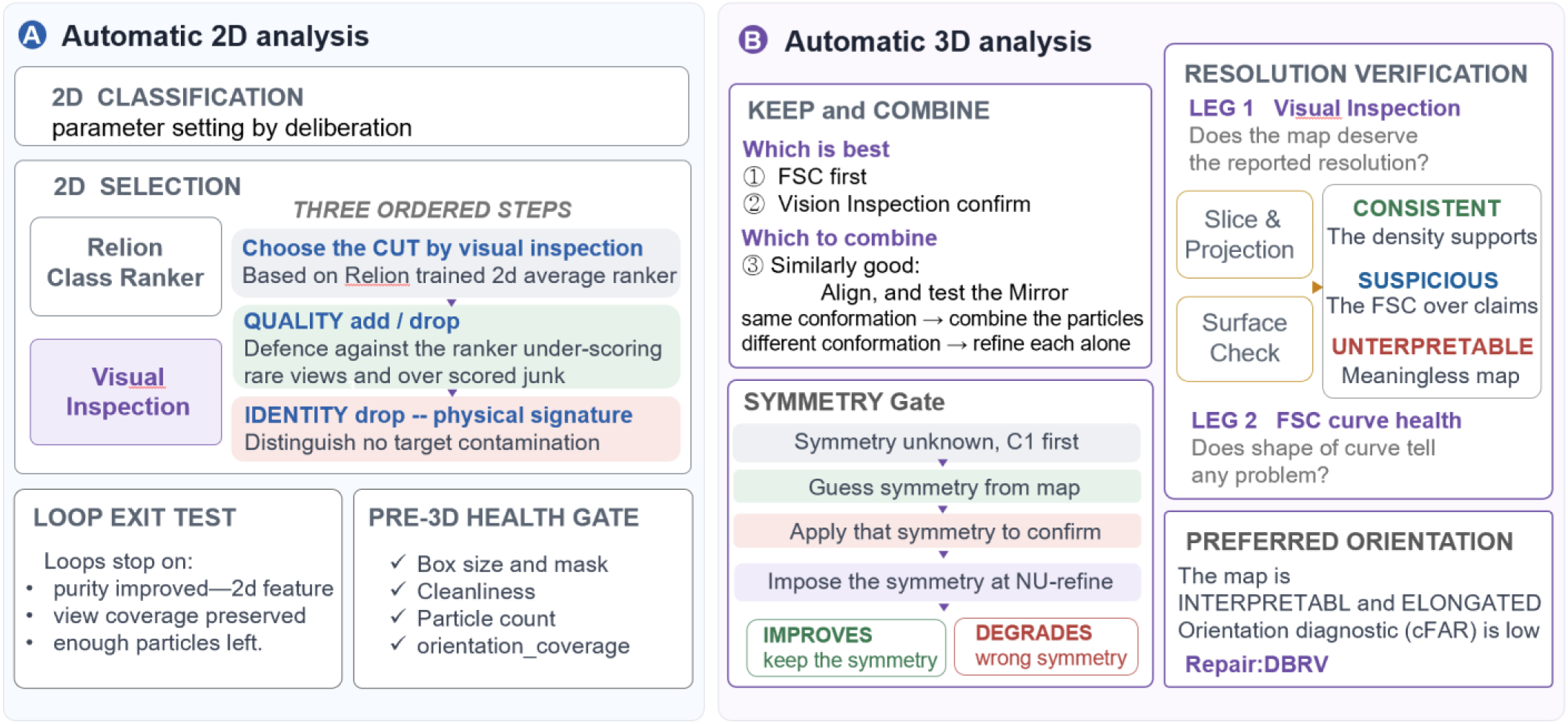
Autonomous judgement layer for two-dimensional and three-dimensional analysis. (A) Two-dimensional class averages are ordered using the RELION class ranker and assessed by image inspection to choose a score cut, correct quality-based omissions or inclusions, and remove classes inconsistent with the target. Loop exit criteria and a pre-3D health gate combine particle geometry, cleanliness, particle number, and orientation coverage. (B) Three-dimensional candidates are compared using GSFSC and image evidence; similar conformations may be aligned and combined, symmetry is tested from C1, and the reported resolution is checked against map slices, projections, surfaces, and FSC-curve health.

##### 2D analysis (Figure 2A)

In EM-Claw, we use RELION class ranker^[7]^ to score the class averages, and the agent inspects a contact sheet ordered by those scores. Selection begins with a score cutoff chosen through the vision-capable LLM’s visual assessment. The agent then adds or removes individual classes on the basis of image quality and can optionally remove classes assigned to off-target proteins by comparing them with the physical description of the specimen (Figure 2A). By combining learned scores with visual assessment and specimen-specific information, this approach helps address the limitations of selection based solely on a pretrained ranking model, allowing class selection to be adapted to the dataset.

The 2D cleaning proceeds as an explicitly counted loop. Before the first round, the exit criteria are recorded: purity must improve, orientation diversity must be preserved, and sufficient particles must remain. The executor runs and evaluates each round, whereas the orchestrator maintains the round counter and audits the stopping decision.

Before the first 3D job of a phase, the particle set is evaluated using four checks: box and mask geometry relative to particle size, cleanliness, particle count, and orientation coverage. Each check is assigned to the instrument that provides the most reliable evidence. Particle size is estimated from the averaged particle images. Cleanliness is assessed visually but interpreted relative to the extraction Nyquist limit, so that a resolution limit caused by binning is not misidentified as contamination. Particle count is advisory and interpreted in light of symmetry. Orientation coverage is determined from the agent’s count of genuinely distinct views. The gate returns one of five verdicts—healthy, minor, more 2d cleaning, serious, or ask human—each of which maps to a predefined orchestrator action.

##### 3D analysis (Figure 2B)

For a 3D reconstruction job that yields multiple volumes, the choice of reconstruction and particles to carry forward is based on measurement. When a job reports a gold-standard FSC^[38]^, the FSC is the primary selection criterion and visual inspection acts as a safeguard. Ab initio reconstructions do not report an FSC in cryosparc 4, in that case, the agent first determines whether a candidate volume resembles protein density and then compares candidates using measurements of volume integrity and continuity. A decision made without an FSC is recorded as provisional and is explicitly confirmed or overturned when a downstream refinement produces an FSC. When candidates are approximately equivalent, the volumes are aligned to one another with mirror-related alignment included. Particles from classes that represent the same conformation are combined, whereas genuinely distinct conformations are refined separately rather than merged.

Symmetry is likewise inferred from the data rather than assumed. When symmetry has not been established, reconstruction proceeds in C1. Candidate point groups are detected from the C1 map and evaluated by symmetrization and overlap. A candidate symmetry is imposed only when its application improves the map, and the improvement is then confirmed.

An FSC-derived resolution estimate can be misleading even when a gold-standard procedure is used^[39, 40]^. EM-Claw therefore combines direct evaluation of the FSC with visual verification of the reconstruction. The FSC is read from the curves stored with the job, and central slices and projections of the map are submitted for visual resolution estimation. When the difference between the two estimates exceeds the predefined tolerance, the system requests a second inspection of the volume surface. The shape and overall health of the FSC curve are also analyzed as secondary evidence to identify potential problems in the reconstruction.

#### 1.6 Experience and self-evolution

EM-Claw retains experience at three levels. First, a per-project record stores every phase, job, and run-time selection together with its numerical basis. It also contains a ledger that labels unproductive branches without deleting them, so that a negative result remains available for deliberate reconsideration. Second, a persistent cross-project memory contains lessons distilled from measurements. Each lesson states the measurement on which it is based and the conditions under which it applies. A lesson is explicitly retracted when later evidence contradicts it, allowing the memory to remain revisable rather than accumulate error. Third, a successful project is distilled into a reusable pipeline notebook. Proven parameters are retained, dimensional parameters are recalculated for the new specimen, and every run-time judgement remains an explicit decision cell supported by the same tested decision procedures used by the agents. The experience accumulates across datasets without model retraining, consistent with experiential and workflow memory developed for language agents^[41–48]^.

#### 1.7 Human interaction

The human can make indications at the beginning of a phase and reviews the reconstruction at its end. That feedback becomes an input to evidence gathering in the next phase. Intervention is not limited to phase boundaries: the human can redirect the run at any point in natural language, after which the orchestrator re-enters the same gather–plan–validate cycle rather than applying an ad hoc change. An ambiguous verdict or failed health check can halt automated processing and request human inspection before further GPU time is committed. Users can also set how independently EM-Claw works through conversation. It can ask for approval at each stage or run on its own.

#### 1.8 Implementation

EM-Claw is implemented in Claude Code^[21]^ and controls CryoSPARC (v4.7)^[1]^ and RELION (v5)^[34, 35]^ through the local Model Context Protocol (MCP)^[49]^ server. The orchestrator (agentM) and executor (subC) run on Claude Opus. The plan validator (subV) runs on GPT through the Codex command-line interface^[22]^, and the remaining specialized agents run on Sonnet. ChimeraX^[50]^ is used for map rendering and surface inspection; model prediction and docking for model-derived masks use the tools described in Section 2.

### 2. Tools

EM-Claw is not a closed pipeline restricted to fixed cryosparc job types. Because the agents write code rather than populate a static configuration, any program with a command-line interface or Python API can be incorporated into a plan. The orchestrator reads the tool documentation and writes the call, and the executor debugs the call against the running system.

Two types of tools enter through this layer. The first comprises published algorithms that outperform a built-in step for a particular problem. For example, EM-Claw can run RELION either within a CryoSPARC workflow or independently of CryoSPARC. CryoRanker is included as an option to rank particles by quality^[10]^, ChimeraX^[50]^ and Phenix^[51]^ are also integrated, allowing the agents to access them as needed. The architecture also allows users to ask the agent to install, configure, and run other software of their choice during a session to meet project-specific needs.

#### RELION integration (Figure 3)

EM-Claw controls RELION (v5)^[34, 35]^ through a local Model Context Protocol (MCP) server^[49]^. The design separates scientific intent from native execution through version-controlled operation definitions, option manifests, parameter mappings, and workflow dependencies. Before submission, the server validates inputs and parameter combinations, translates them into relion_pipeliner jobs, and registers each job in the native pipeline. EM-Claw then plans, monitors, and evaluates the workflow through a bounded plan–execute–assess–decide cycle. This separation preserves RELION’s job organization, outputs, and provenance while keeping agent actions inspectable and reproducible; successful execution remains distinct from scientific acceptance. The integration supports movie import, motion correction, CTF estimation, particle picking and extraction, 2D and 3D classification, initial-model generation, refinement, masking, postprocessing, CTF refinement, and Bayesian polishing. An optional bridge uses pyem^[52]^ to transfer selected CryoSPARC particle metadata and image stacks into RELION for further processing.

**Figure 3.**
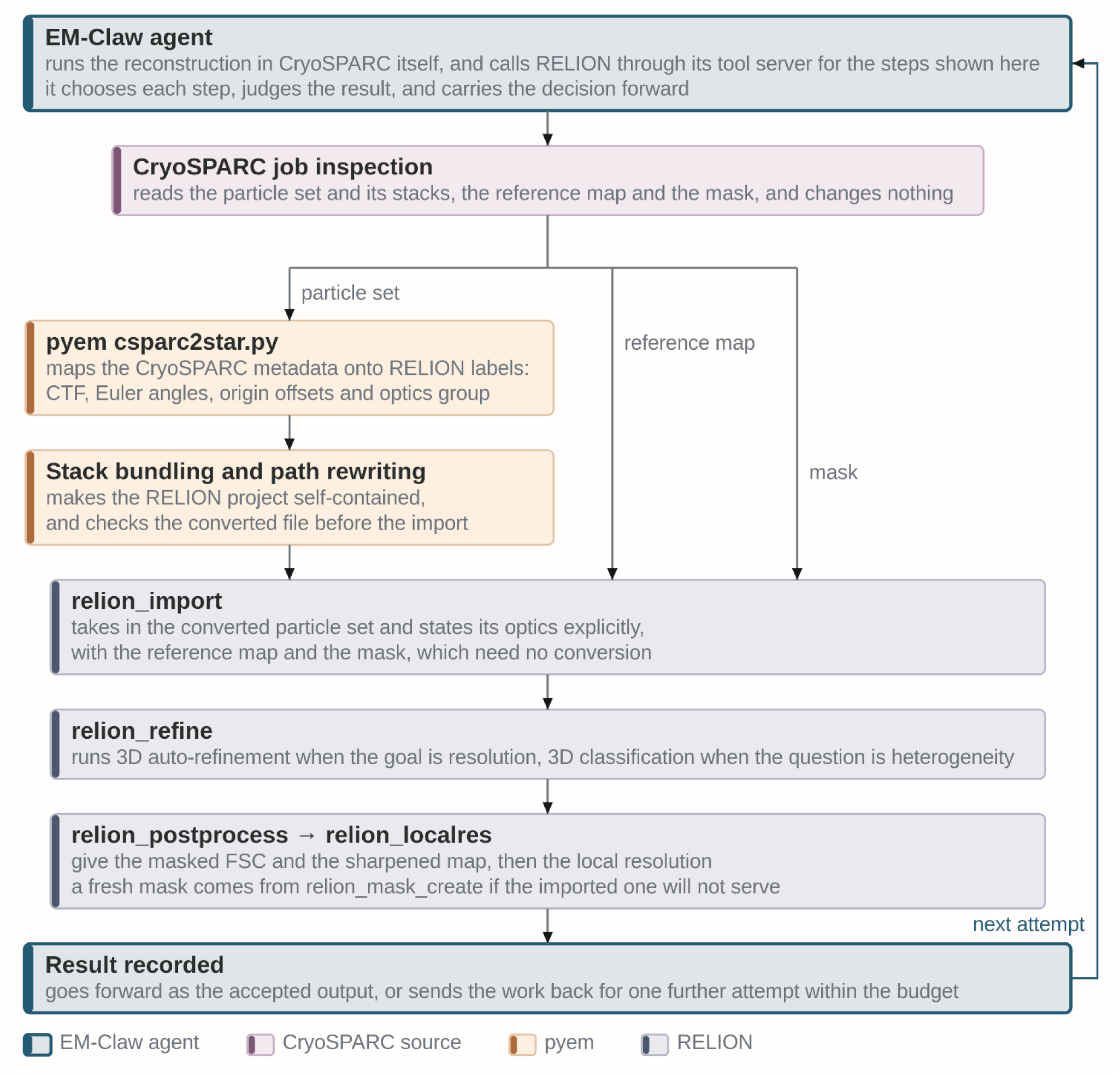
Transfer of CryoSPARC particles to RELION through pyem. The particle metadata needs conversion: pyem’s csparc2star.py maps the CTF parameters, Euler angles, origin offsets and optics groups onto RELION labels. The image stacks are bundled into the RELION project with their paths rewritten, and the converted file is checked before import.

The second type comprises purpose-built tools that encode user experience for a problem presented by a particular specimen. Two such tools included in EM-Claw are described below (Figure 4), users can also develop their tools inside the agent.

**Figure 4.**
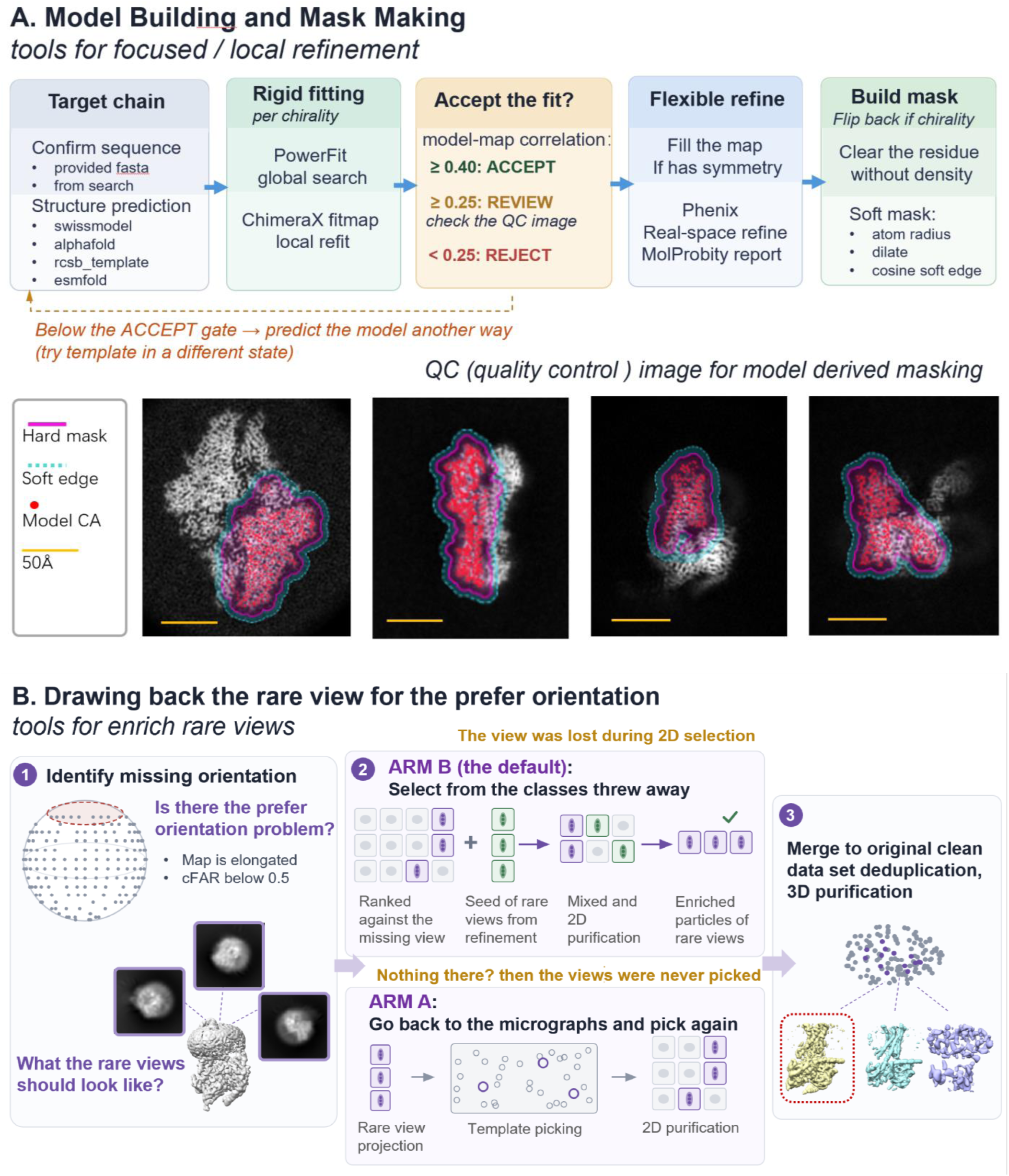
Examples of purpose-built tools that encode user experience. (A) Tools for building model and making mask. (B) Tools for gathering more particles in rare views.

#### Model-derived masks for local refinement (Figure 4A)

Focused refinement requires a mask confined to the region of interest. A density threshold is sufficient when that region can be separated by density alone, but it cannot isolate one chain, one domain, or a ligand-binding pocket from the remainder of a complex. Therefore, to generate precisely localized masks, we developed a tool for constructing masks from atomic models built in EM-Claw.

The sequence is first confirmed from the fasta file recorded for the project or, if no sequence file is available, by searching an external source. A model is then predicted and docked into the map by a global search followed by local refitting, with both hands tested. The fit is evaluated using the model-to-map correlation: a value greater than 0.40 is accepted, a value from 0.25 to 0.40 is reviewed against the rendered quality-control image, and a value below 0.25 is rejected. If the fit fails this gate, the procedure returns to model prediction or template selection rather than generating a mask from a model unsupported by the density. The conformational state of a template is supplied explicitly because a template in the wrong state can dock poorly even when the overall fold is predicted correct. After rigid-body fitting, optional flexible fitting with Phenix can be used to improve the model. A radius is then applied around the selected atoms, the resulting region is dilated, and a cosine soft edge is added with a width scaled to the map resolution. Before use, the mask is inspected in a rendered slab view. A mask volume between two and three times the protein volume is considered acceptable.

Because the mask is defined by the model, the target can be specified in sequence terms, including selected chains and residue ranges, rather than by a contour level. A region defined by the biological question rather than by density contrast can therefore be isolated.

#### Recovery of rare views under preferred orientation (Figure 4B)

When a reconstruction is limited by preferred orientation, particles representing the missing views may be absent from the processed particle set rather than from the specimen itself, they may have been lost during the picking, or during 2D selection. Drawing back rare views (DBRV) is designed for such situations after a preliminary 3D refinement when the overall 3D information for the sample is available.

The orientation distribution of the refined map is examined first, and the map is projected along the under-represented directions to define the target views. Two routes then return to the particles. In the picking route, these projections are used as templates for a new pass over the micrographs, recovering views that may not have been picked initially and enriching them sufficiently to support grouping. In the selection route, class averages rejected in every round and branch are ranked against the under-represented projections. Particles from the matched classes are combined with a seed set of particles that the refinement assigned to the deficient directions and are then reclassified. The seed helps the few genuine rare-view particles to group against dispersing among the rejected particles. In either route, the recovered particles are 2D purified before being merged with the clean particle set, with duplicates removed.

## Results

Before applying EM-Claw, a digested precedent bank needs to be built based on individual lab’s processing experience. A deterministic script scanned every project directory in our lab’s shared CryoSPARC storage. Within each project, it grouped jobs into independent processing lineages and generated a summary for each lineage if the processing reached resolution under 4 Å. Across 176 project directories associated with 51 users, the scan identified 463 qualifying lineages. Then the agent digested the raw history into its precedent databank. A selected collection of six processing workflows from experienced laboratory members was manually curated, given with higher priority during retrieval. These workflows cover diverse specimen types and processing strategies, including two GPCR samples, one transporter monomer, one soluble protein sample with prefer-orientation, one small soluble protein of 50kDa in C2, one membrane protein complex in C4.

We first selected four publicly available cryo-EM datasets: β-galactosidase (RELION tutorial subset of EMPIAR-10204^[53]^), influenza haemagglutinin trimer (EMPIAR-10097, 40° tilt; EMPIAR-10096, untilted^[11, 54]^), and TRPV1(EMPIAR-10059^[54, 55]^)—to systematically evaluate EM-Claw. These datasets encompass proteins with diverse symmetry properties and include both soluble and membrane proteins, providing a representative benchmark for assessing EM-Claw’s ability to perform general-purpose, end-to-end single-particle analysis.

We then applied EM-Claw to recent or ongoing research projects in our laboratory to demonstrate its practical applicability, where the sample appears in lower quality and more difficult for processing (Figure S10), reflecting the challenges encountered in routine single-particle analysis.

### 1. Evaluations on common cases from public EM data bank (Figure 5)

#### β-galactosidase dataset (Figure 5A, S1)

The RELION β-galactosidase tutorial dataset^[53]^ (24 movies) is compact and suitable for initial testing, with the main challenge being to recover enough usable particles from limited data. We processed it using three independent EM-Claw instances, which adopted different strategies for particle recovery and cleaning.

**Figure 5.**
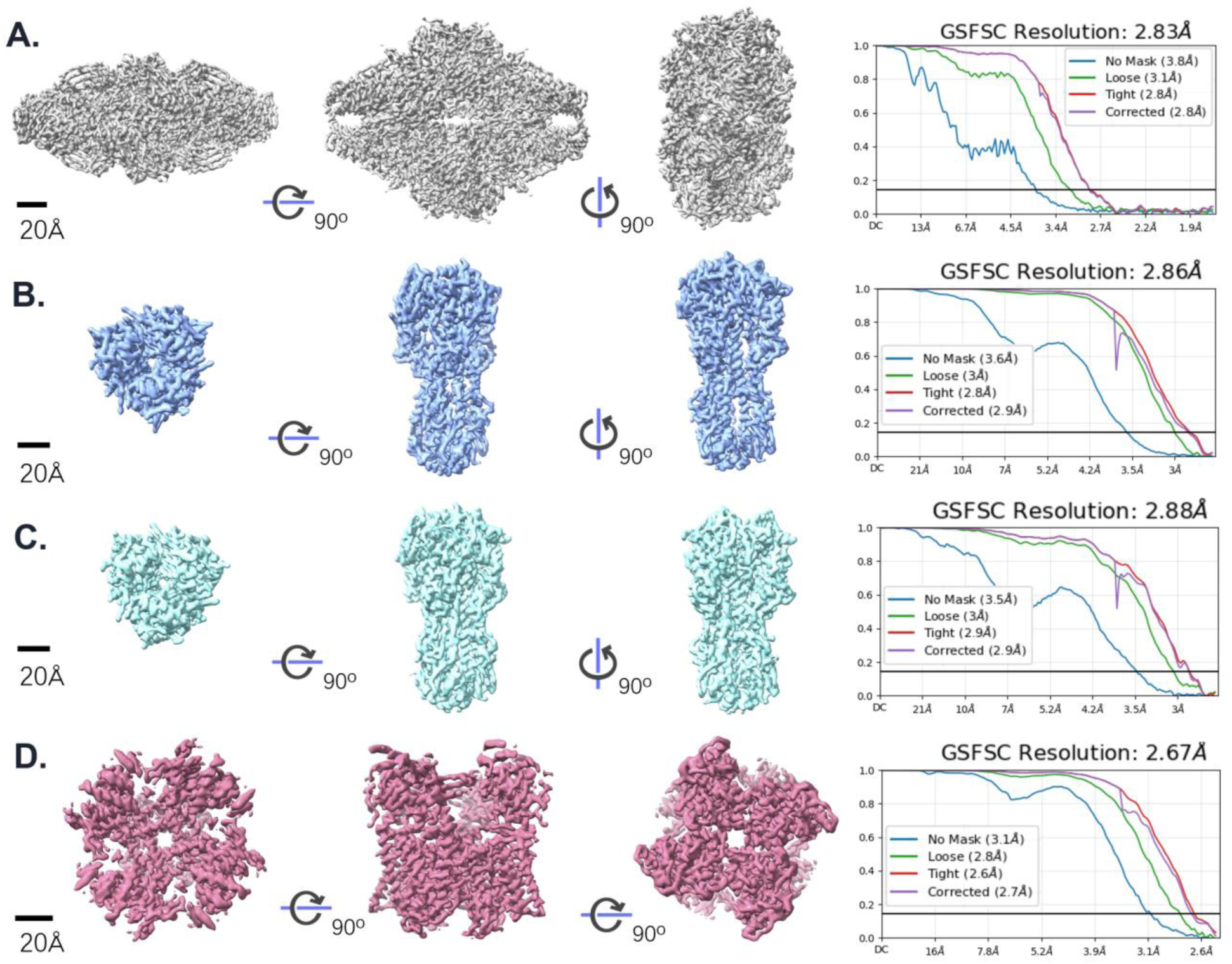
Reconstructions of four public benchmark datasets by EM-Claw. Each row shows three orthogonal views of the final map and its gold-standard FSC curves. (A) β-galactosidase from the RELION tutorial dataset, D2, 2.83 Å, compared with tutorial’s 2.97 Å. (B) Haemagglutinin trimer from data collected at a ∼40° stage tilt (EMPIAR-10097), C3, 2.86 Å, compared with published 4.2 Å(EMD-8731). (C) Haemagglutinin trimer from untilted data with severe preferred orientation (EMPIAR-10096), C3, 2.88 Å, compared with the published failed reconstruction. (D) TRPV1 in nanodiscs (EMPIAR-10059), C4, 2.67 Å, compared with published 2.95 Å (EMD-8117, PDB 5IRX).

The processing started from raw movies, with no reference model and no prior assumption of symmetry. The agent worked out the gain-reference orientation, the extraction box size itself and confirmed D2 symmetry from the reconstructed map.

The first instance (Figure S1A) briefly combined blob, template, and Topaz picking^[4]^, removed duplicate picks, and retained 4,706 particles after one round of lenient 2D selection. A subsequent two-class ab initio reconstruction identified a subset of 3,383 particles, which yielded a 3.02 Å reconstruction.

The second instance (Figure S1B) used blob picking followed by three rounds of template re-picking. It maintained both a 2D-selected pool of 6,426 particles and an unfiltered pool of 12,864 particles. Heterogeneous refinement of the unfiltered pool, using decoy maps generated from rejected particles, recovered 1,898 particles excluded during 2D selection. It also found extensive overlap between picking methods: 98.2% for blob and template picks and 99.5–99.8% for Topaz picks, indicating little scope for further particle recovery through additional picking. The final set of 6,448 particles yielded a 2.83 Å reconstruction (Figure 5A).

A third run (Figure S1C) tested interoperability between CryoSPARC and RELION. EM-Claw transferred particles from CryoSPARC to RELION for 3D classification, then returned the selected particles to CryoSPARC for refinement. This workflow successfully produced a 2.86 Å reconstruction from 5,473 particles.

These independent runs illustrate the flexibility of LLM-driven processing: the agent can devise different routes to address the problem which is the key to enable further improvement.

#### Influenza HA trimer (Figure 5B and C, S2 and S3)

We next applied EM-Claw to 668 motion-corrected micrographs of the influenza HA trimer, a specimen known for strong preferred orientation, starting without a reference model. Without being told how the data were collected, the agent found from per-particle defocus gradients that the entire dataset had been recorded at a ∼40° stage tilt. Two independent internal checks confirmed this, and it matches the published collection^[11]^. The three-fold symmetry was already visible in the C1 reconstruction (3.17 Å), and imposing C3 improved the map to 3.05 Å. In a second round, the agent tested two ways to improve the map separately. CTF refinement alone improved the map by 0.07 Å, to 2.98 Å. Particle enrichment by template and Topaz re-picking (+48.6% particles) gave 3.02 Å, a gain of only 0.03 Å. Refining the CTF across the whole enlarged particle set then gave 2.89 Å, and local refinement gave the final 2.86 Å C3 map from 303,305 particles. Its visually estimated resolution agreed with the FSC value, and it showed a resolved α-helical stem and separated head protomers (Figure 5B and S2).

As a harder test, we processed the untilted HA dataset, which is known for severe preferred orientation, from 447 motion-corrected micrographs without a reference model. The agent ran blob, template and Topaz picking in parallel and judged each picker by the views it added, not by particle count. Template picking mostly added more top views, whereas Topaz recovered side views, so the agent built the final particle set from blob and Topaz picks only and left the template picks out. The agent also noticed that the second round of heterogeneous refinement removed side views preferentially. A control refinement that kept all 111,954 particles matched the cleaned set in resolution (2.89 versus 2.92 Å) and had better angular coverage, so the agent dropped that second cleanup round. Per-particle defocus and global CTF refinement then gave the final C3 map at 2.88 Å (3.47 Å without a mask), showing resolved α-helices and β-strands (Figure 5C and S3).

#### TRPV1 channel (Figure 5D and S4)

We next processed 1,200 motion-corrected micrographs of the TRPV1 channel in nanodiscs, starting without a reference model. From two rounds of 2D classes it took a strict, view-balanced subset to build the 3D references and a looser subset as the particle pool. Two rounds of heterogeneous refinement against those references left 207,017 particles. Before adopting C4, the agent compared it with a matched C1 refinement using an aligned symmetry axis; C4 improved resolution and gave a cleaner FSC curve. The final C4 map reached 2.67 Å (Figure 5D), compared with the published resolution of 2.95 Å for the same dataset (PDB 5IRX; EMD-8117)^[55]^.

### 2. Towards higher resolution: Two actual lab cases of GPCR reconstruction with clear ligand density recovered (GCGR and SSTR5)

#### GCGR (Figure 6 and S5)

To evaluate EM-Claw in practical research situations, we reprocessed two datasets from recently published studies in our laboratory: a GCGR–Gs complex^[56]^ and an SSTR5 dimer with Fab^[57]^.

**Figure 6.**
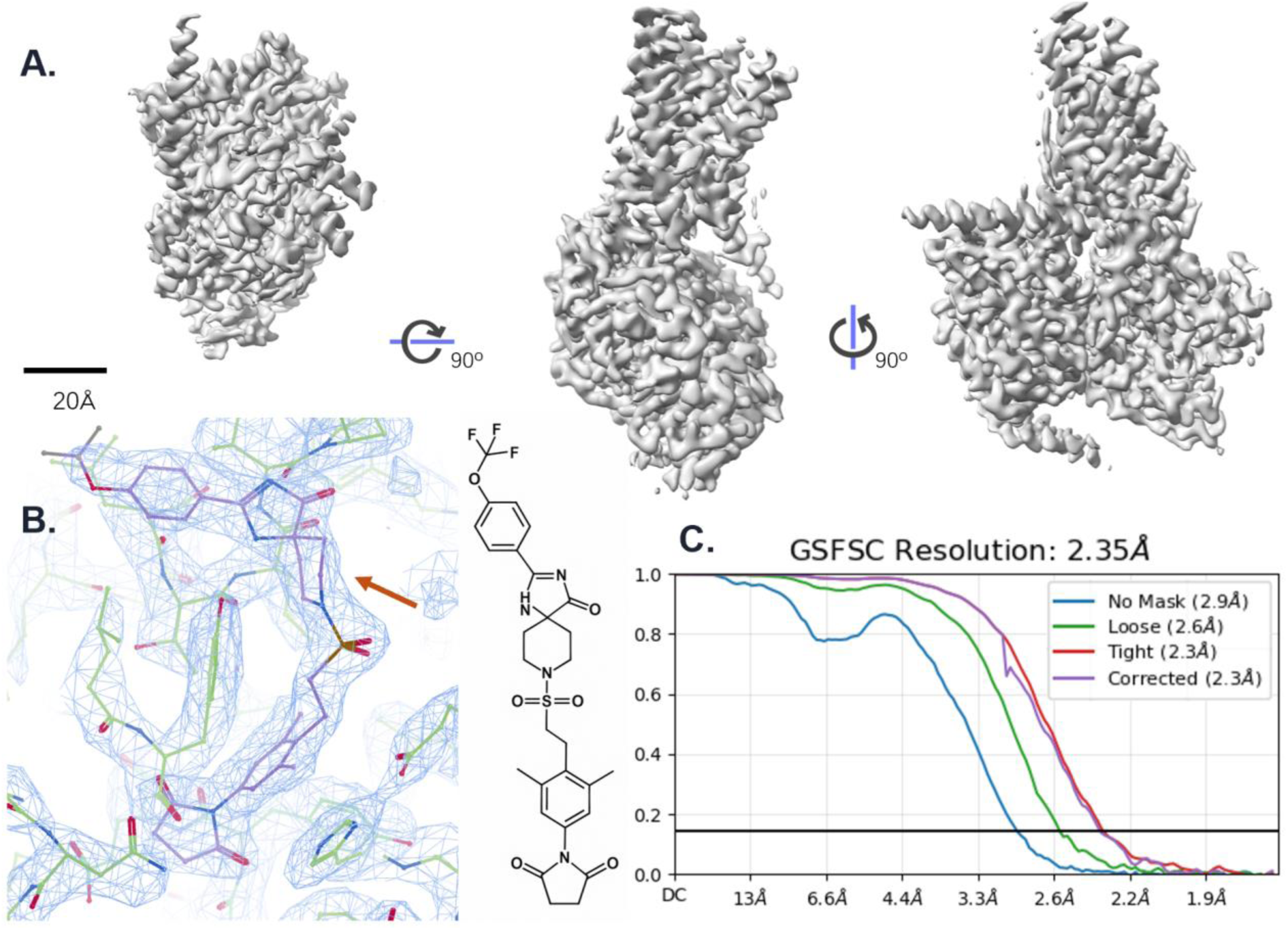
(A) Global map of GCGR–Gs with ligand in allosteric pocket reconstructed by EM-Claw (without sharpening) and (C) the corresponding FSC. (B) With improvement of the resolution to 2.35 Å compared with to the published 2.53 Å (PDB 9XN4, EMD-67038), the chair conformation of a six-membered ring (red arrow) was confirmed in the sharpened map, which was figured only by binding energy for analogous compounds previously reported by our group.

For the GCGR–Gs complex, EM-Claw processed 5,991 motion-corrected micrographs without access to previous processing of this dataset, drawing instead on acquisition metadata and precedents from other datasets. Particle picking, two rounds of 2D classification, and 3D classification yielded 264,383 particles, producing a 2.68 Å reconstruction. The agent then systematically evaluated potential improvements, identifying beam-tilt and trefoil correction as the main contributors to a final resolution of 2.35 Å (Figure 6A and C), compared with the previously reported resolution of 2.53 Å for the same dataset (PDB 9XN4; EMD-67038)^[56]^. The improved map clearly resolved the transmembrane helices, the Gβ propeller blades, and the allosteric ligand, lending stronger support to the ligand’s binding conformation (Figure 6B).

#### SSTR5 (Figure 7 and S6)

We next processed 4,619 motion-corrected micrographs of antagonist-bound SSTR5. The agent noticed that every 2D round removed a large part of the particles without converging and suspected that the selection was discarding real but blurred particles. The initial consensus reconstruction reached a global resolution of 3.68 Å but exhibited strong anisotropy, with directional resolutions ranging from 3.0 to 8.7 Å (cFAR = 0.05). Subsequent heterogeneous refinement of the particles rejected during 2D classification using the consensus map, junk volumes and a decoy built from empty micelles as references, recovered about 64,000 particles and the missing views. A second cycle, with a Topaz picker retrained on the refined particles, the cFAR was finally raised to 0.38 and the resolution reached the sampling limit of the binned data (2.99 Å, 203,713 particles), so the agent re-extracted at the original pixel size. The agent analyzed the result and found the orientation of 19% of the particles flipped by more than 150°. 3D classification could not separate these particles, but the refinement with orientation-stable particles gave a clearly better map than an equal-sized random subset at matched refinement depth. The agent therefore kept only the particles with stable orientation for consensus refinement and local refinement, achieving resolutions of 2.94 Å for the overall complex (Figure 7A and B) and 2.41 Å for the receptor core. Meanwhile, the agent informed the operator to substantial instability in particle orientations, with angular deviations approaching 180°. Upon inspection of the reconstructed density map, the operator hypothesized that conformational heterogeneity of the Fab obscured the intrinsic C2 symmetry of the receptor dimer and requested focused refinement of the receptor region with C2 symmetry imposed. Then the agent transformed the alignment parameters into the C2 symmetry frame and performed symmetry-constrained local refinement, dramatically improving the resolution of the receptor to 2.16 Å (Figure 7C, D and E), compared with 2.59 Å in the previously published reconstruction (PDB 9W32; EMD-65588)^[57]^.

**Figure 7.**
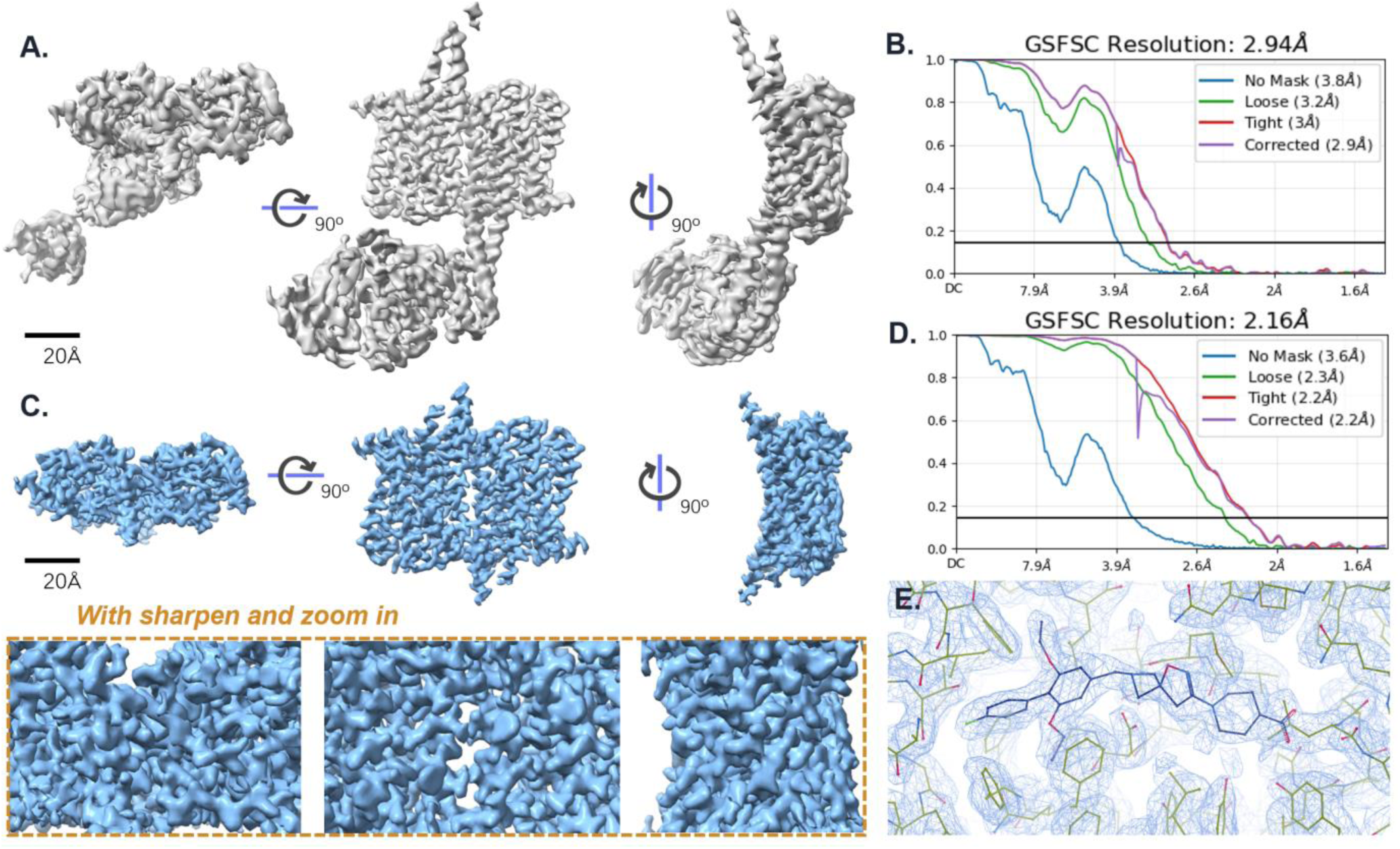
(A) Global map (without sharpening) of the antagonist 1–SSTR5 complex and (B) the corresponding FSC curve. (C) Local refinement focusing on the receptor with applying C2 symmetry recovered the very detail of the sidechain’s structure with the resolution of 2.16 Å with FSC showed in (D), compared to the published 2.59 Å (PDB 9W32, EMD-65588). The upper raw: orthogonal views of the map without sharpening, the lower row: corresponding sharpened map with zoomed in views in the center part. (E) The improvement of the resolution provides solid information of the bind pose and interaction for the ligand in the pocket.

At this step, the agent strongly suggested using original movies to push the resolution further to break the resolution record for EM structure of GPCR, but unfortunately the movies of the dataset have already been deleted due to storage limitations.

### 3. Reconstruction for heterogeneous and unstable sample: a hard lab case for CB1 complex

We then tested with 8,326 micrographs of human cannabinoid receptor 1 (CB1) bound by an Fab antagonist (Figure 8, S7). The dataset’s difficulty lies in its composition: intact complexes (the target particles) are present amid a large amount of dissociated debris (Figure S10G).

**Figure 8.**
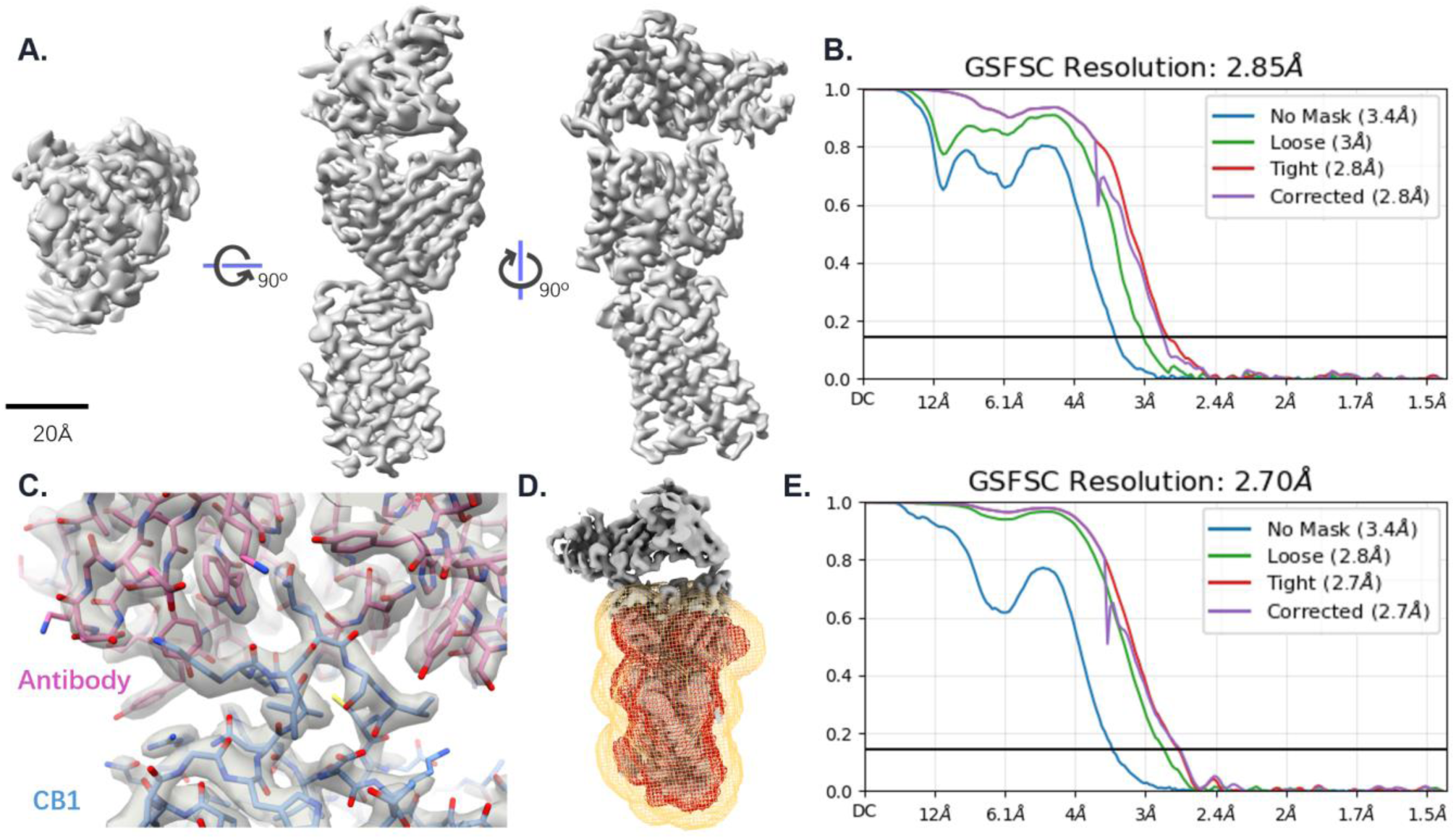
(A) Global reconstruction of CB1–Fab by EM-Claw (without sharpening) and (B) the corresponding FSC. (C) The sharpened map with the built model of the Fab interaction area. Local refinement with the model derived soft mask precisely covering receptor and the interface of Fv improved the resolution of receptor and the interaction interface (E). (D) The mask used in the local refinement, red: mask of density value 1 and orange: the soft edge.

The first campaign failed and the operator requested for a fresh trying with identifying what was happening: the inactive-state specimen is unstable, so the Fab-bound complex is a minority population mixed with receptors that carry no Fab, alongside empty-micelle look-alikes. The second campaign was run in a fresh session with the lesson: “Target of this phase: the antibody-bound complex, not the bare receptor.” The small group of complex-shaped class averages showing the connected two-lobe morphology were selected and reconstructed ab initio (Figure S10G). The best class gave a 5 Å reconstruction of the receptor-Fab complex, which was used for generating projection templates for further particle picking.

The decisive step was the 3D classification that followed. After a single round of removing obvious junk particles by 2D selection, the agent decided to sort the full, species composition-mixed pool with hetero refinement against five references: one positive reconstruction of the complex and four decoy ones built from the dataset. The strategy reduces the loss of good particles when the amount of interfering particles is large, and the resulting best class yielded a 2.93 Å reconstruction. The agent renewed its positive reference to perform another hetero refinement, and the best class was refined to 2.90 Å. Then the operator asked to merge the particles from the two refinements, and the agent decided to also try the intersection of the two datasets, which gave a higher resolution than the merged set: 2.85 Å (Figure 8A, B).

In phase 3, the operator named the scientific objective: the receptor–Fab interaction is what the study is about. The agent addressed it with focused refinement using a mask derived from modelling, covering the whole receptor and the interface area of the Fv (Figure 8D), and the resolution of the focused area was improved to 2.70 Å (Figure 8E), with side chains at the interaction interface clearly seen (Figure 8C). The method for making the mask has been described in the Methods section. The result of the agent’s solved structure is on submission in parallel with another paper of our lab^[58]^.

### 4. Brief solutions for small membrane sample: Transporter application cases with reusable pipeline

Transporters are particularly challenging specimens for single-particle cryo-EM. They are small, have limited extramembrane density to support alignment, and are surrounded by a large, largely featureless detergent micelle. In the specimens studied here, the ordered protein core is approximately 50–60 kDa within an overall particle envelope of about 100 Å. The micelle therefore occupies much of the particle volume without contributing interpretable protein features. We applied EM-Claw to two unpublished transporter datasets collected in our laboratory (Figure 9 and S8, S9).

**Figure 9.**
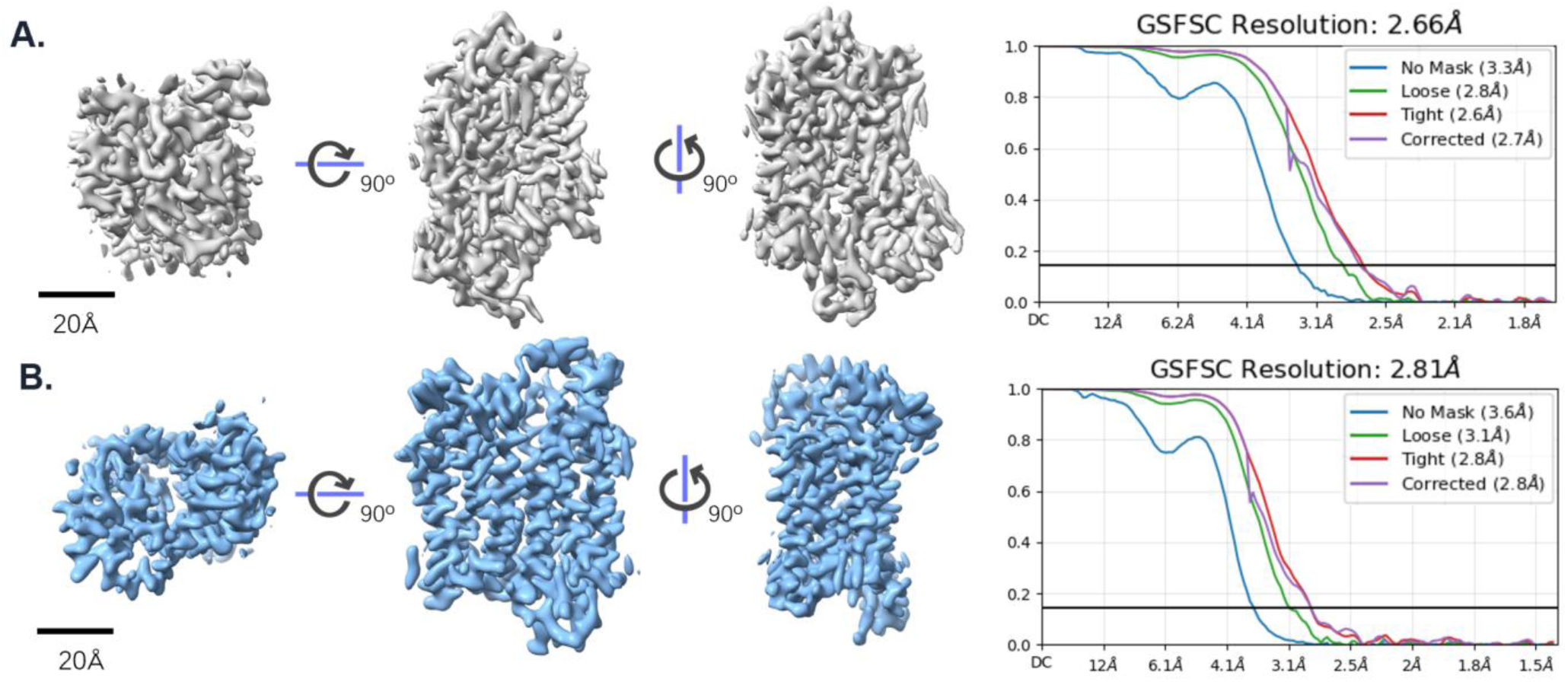
Two different kinds of transporters reconstructed by EM-Claw and the corresponding FSC, for unbiased judgment, all maps displayed are not sharpened. (A) transporter A, local refinement on its protein density: 2.66 Å; (B) transporter B with the reusable pipeline, local refinement on its protein density: 2.81 Å.

Before planning the processing of transporter A, subR retrieved three precedents from the digested history bank. Two were independent processing runs of the same specimen by different human experts; the third involved an unrelated transporter. The operator allowed the agent to proceed without intervention to assess how it would act from these different solutions. EM-Claw drew on the processing strategy of the first precedent and parameter settings from the other two. Compared with the human expert runs, EM-Claw applied less stringent 2D class selection, retaining diverse particle views even when some class averages appeared blurred. For ab initio reconstruction, it used a nondefault combination of an initial resolution of 20 Å, a maximum resolution of 4 Å, initial and final minibatch sizes of 300 and 1,000, respectively, and a convergence epsilon of 0.005, then proceeded directly to refinement. This shorter processing route yielded a 2.66 Å reconstruction of transporter A using 44 jobs (Figure 9A, S8), compared with 175 jobs for a 2.66 Å reconstruction and 115 jobs for a 2.63 Å reconstruction in the two expert runs.

We then asked subP to distil the successful route into an automated pipeline for similar specimens. It identified the jobs contributing to the final map, consolidated the exploratory branches into a single path, and retained the successful parameter settings. Each step requiring a run-time judgement was represented as an explicit decision cell using the same tested decision routines available to the agents. The resulting self-contained Jupyter notebook can run independently of an agent.

The pipeline was subsequently tested on transporter B, producing a 2.81 Å map from 8,567 micrographs (Figure 9B, S9).

## Discussion

Here, we present EM-Claw, a flexible agent system built on Claude Code that supports fully automated cryo-EM single-particle reconstruction. Beyond only benchmarking on public datasets as other similar works, we emphasize capturing processing experience accumulated within the laboratory and applying it to new collected data, reflecting the challenges and working practices of actual structural biology research. The evaluations on both public and laboratory datasets demonstrate its practical value in cryo-EM SPA processing (Table S1).

### Flexible processing routes tailored to working on cases

When cryo-EM specimen quality is high, the choice of processing route may be less critical to the outcome. But many of the structures that remain unsolved are not of that kind. They are the ones that are hard to purify and give poor-quality grids such as the large complexes that fall apart, and the small proteins whose signal-to-noise ratio leaves little to align on. For these, a fixed general-purpose pipeline cannot be right, because the obstacle itself differs from specimen to specimen.

EM-Claw is built on that observation. It composes the route from two things — the properties of the specimen as input and measured from the data, and what has worked before on comparable cases — and it re-decides at each junction rather than following a plan laid down in advance. Similarly to human problem-solving, the same dataset may be processed through different routes: we ran the β-galactosidase dataset three times independently and each time it acted differently. That flexibility is not a by-product, it is the mechanism by which the system gets past a hard case, and the source of whatever improvement it accumulates. In the CB1 case, the first campaign failed but the second succeeded because the agent changed the shape of the route based on the reason of failure. A fixed pipeline has no move of that kind available.

The side effect of flexibility is instability. EM-Claw addresses this at three levels. The first operates on individual decisions. A run that succeeds is added to the history bank, and the runs judged most successful can additionally be placed on a curated short list that the retrieval agent reads before it scans anything else. The second is that, if a route has proven itself, with subP, it can be condensed into a relatively fixed, self-contained notebook for that class of specimen. The third is the substrate beneath both: the persistent memory of the agent framework itself. Lessons distilled from working process are written to files that survive the loss of any single agent’s context and are loaded at the start of every session.

Overall, EM-Claw is designed for the working pattern of most structural biology laboratories, which concentrate on a handful of related proteins rather than arbitrary targets.

### Automation is the goal, the human in the loop is not its opposite

Full autonomy is the natural endpoint of this line of work, and it should not be pursued by refusing help that is cheap and decisive. An EM expert can give the agent valuable suggestions only with a few sentences in natural language, as we did in the case of SSTR5, and the effect is out of all proportion to the cost: current LLM models their multimodal variants do not reason well about protein three-dimensional density and nobody knows a specimen better than the person who prepared it.

For an operator new to cryo-EM processing, EM-Claw can also be used to access the accumulated practice of colleagues and the primary literature, and analyze the experience behind successful history works. The agent supplies breadth and consistency; the operator supplies specimen knowledge and a reliable interpretation of a three-dimensional map.

### Toward an open database of processing experience

Of the system’s inputs, retrieved precedent is the one whose absence is significant. At a very early stage, we built the system with no history-retrieval module. Its reconstructions were markedly weaker, and the plans themselves tended toward the generic — close to a published tutorial workflow and missing steps that turn out to matter on difficult specimens. The reason beneath that may be the lack of a public record of EM data-processing workflows and experience. We would therefore argue for an open, structured database of real single-particle processing trajectories alongside the published structures, which could do more for the automation of this field than further improvements to any individual algorithm.

## Code availability

https://github.com/ericxu-lab/em-claw/

## Author contribution

H.E.X. designed the research; H.S. and J.L. designed and implemented the agent based on claude code. Z.X. provided technical expertise. J.L., Z.X., and S.X. tested the agent. K.W., Q.Y., H.L., L.Z., and Z.X. provided datasets for testing. H.S. and J.L. wrote the manuscript.

---

**Supplementary Figure S1A.**
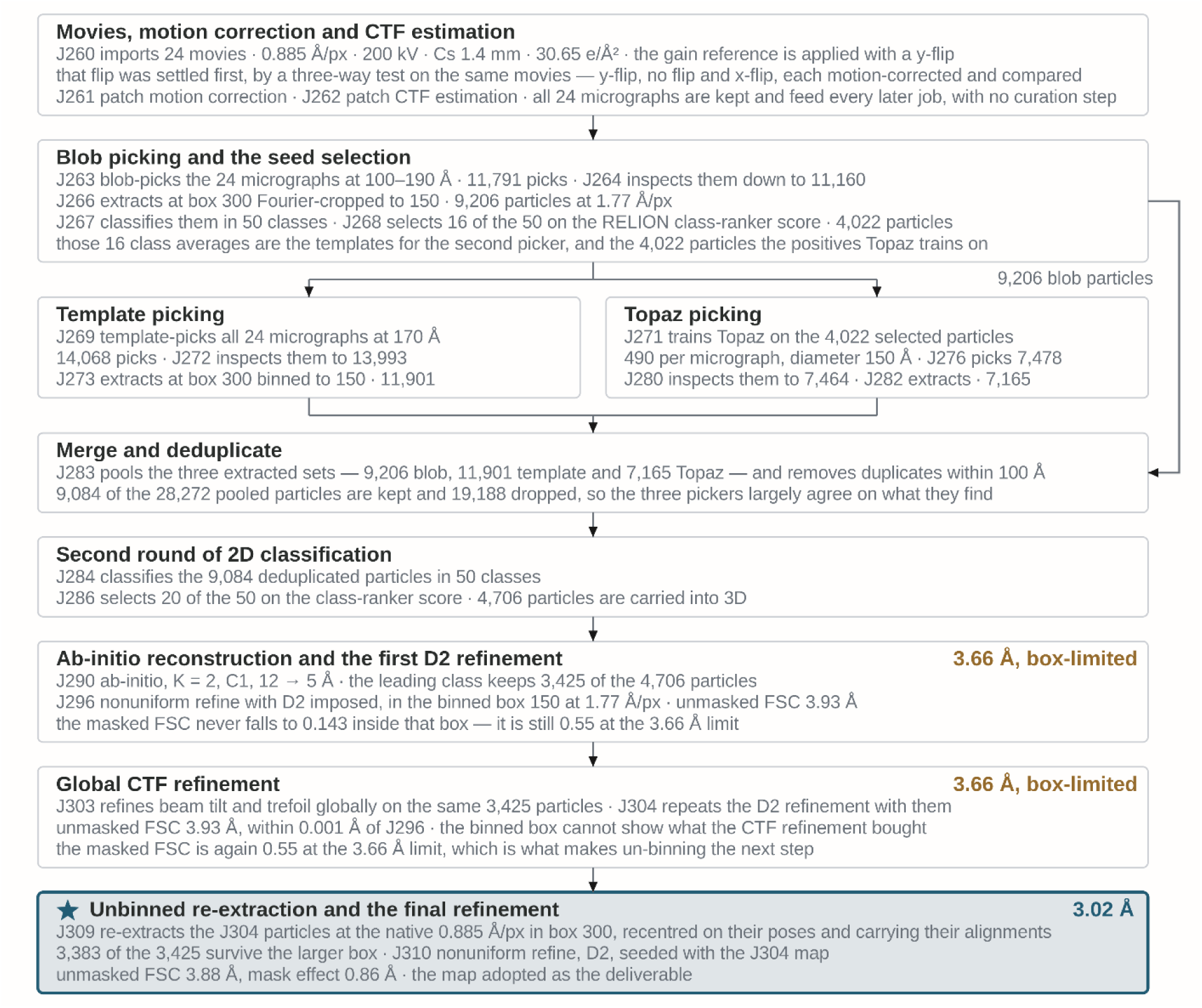
Processing lineage of the β-galactosidase reconstruction (the 1^st^ instance).

**Supplementary Figure S1B.**
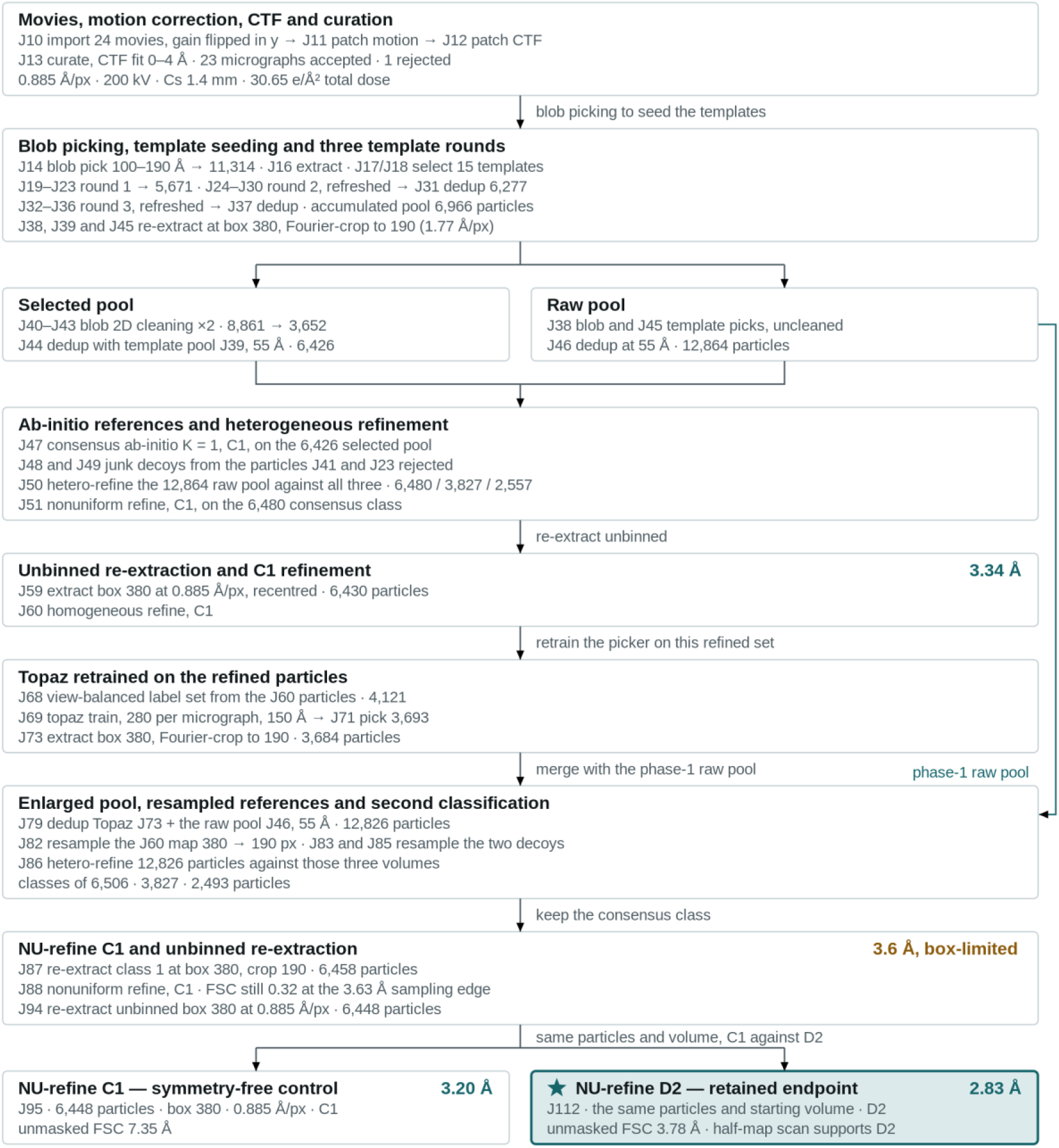
Processing lineage of the β-galactosidase reconstruction (the 2^nd^ instance).

**Supplementary Figure S1C.**
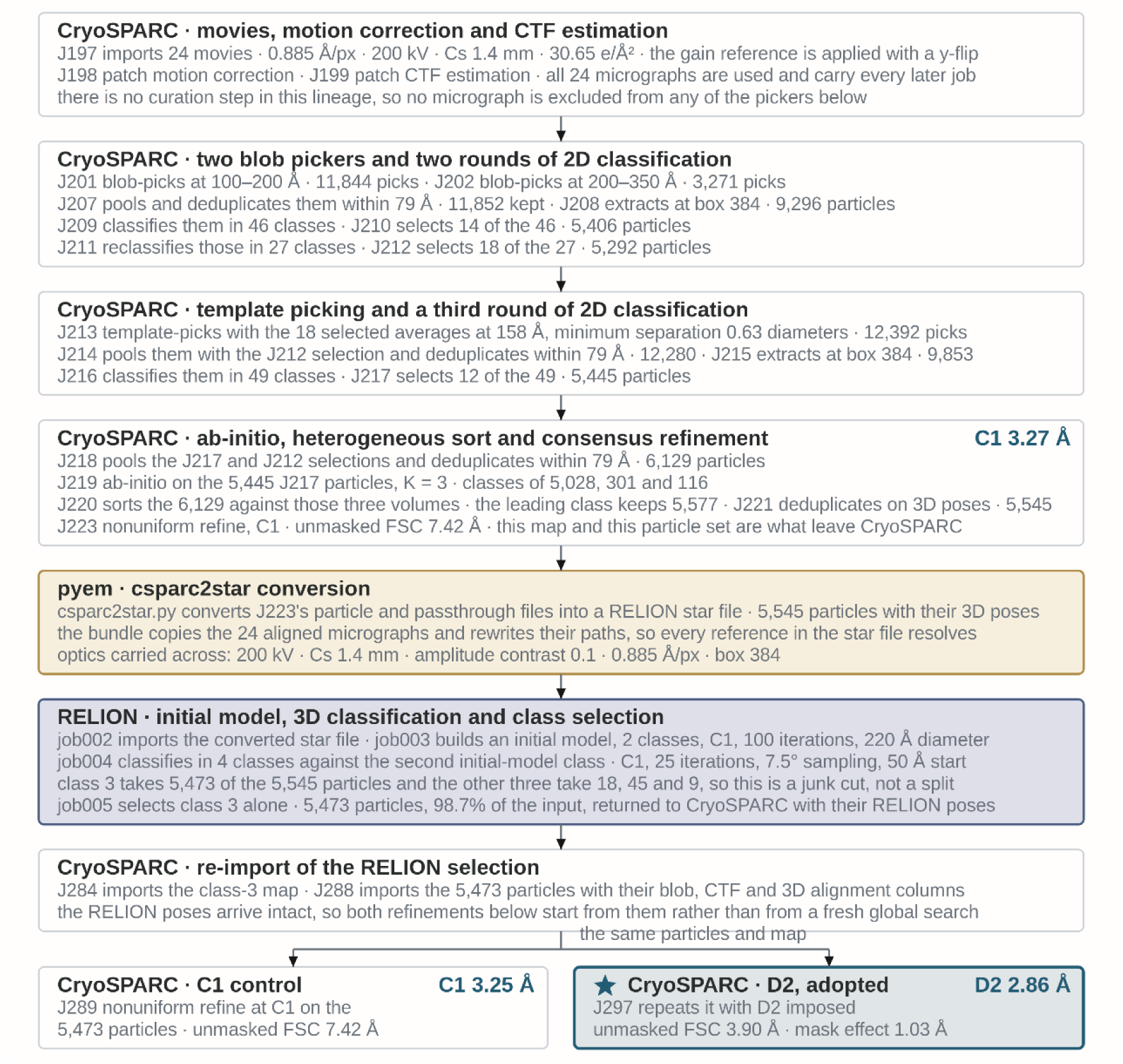
Processing lineage of the β-galactosidase reconstruction (the 3rd instance).

**Supplementary Figure S2.**
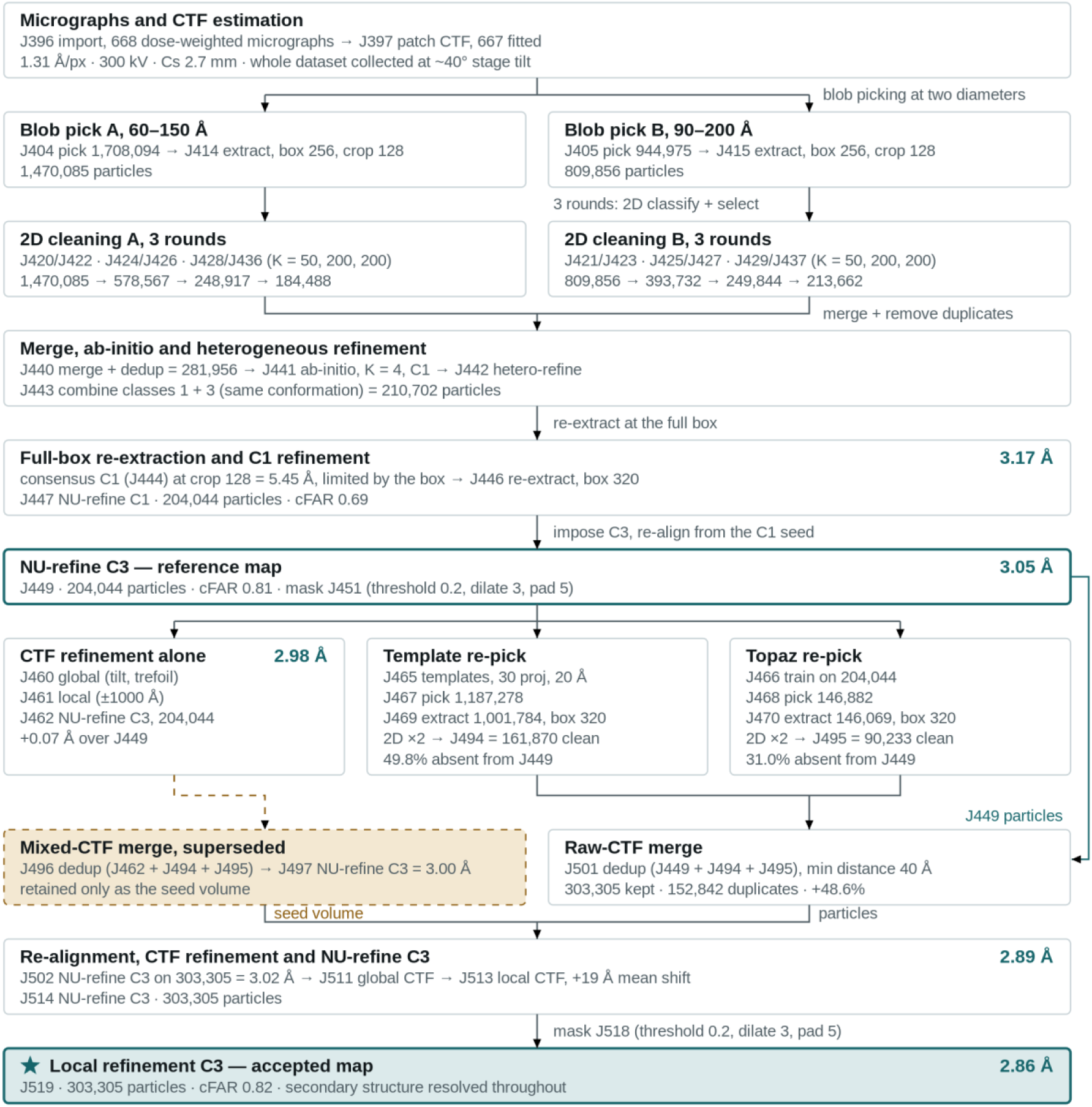
Processing lineage of the tilted haemagglutinin trimer reconstruction (EMPIAR-10097).

**Supplementary Figure S3.**
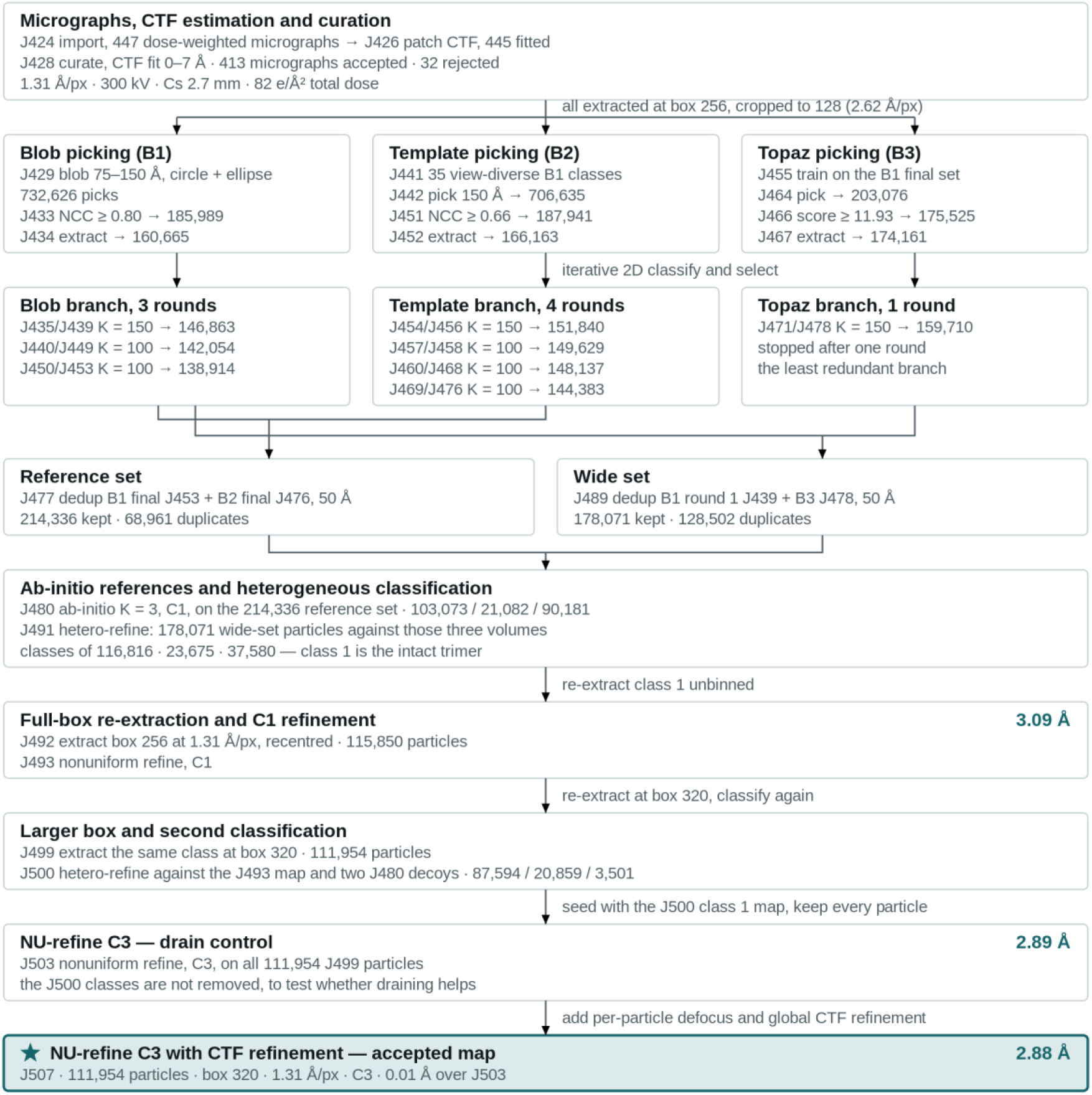
Processing lineage of the untilted haemagglutinin trimer reconstruction (EMPIAR-10096).

**Supplementary Figure S4.**
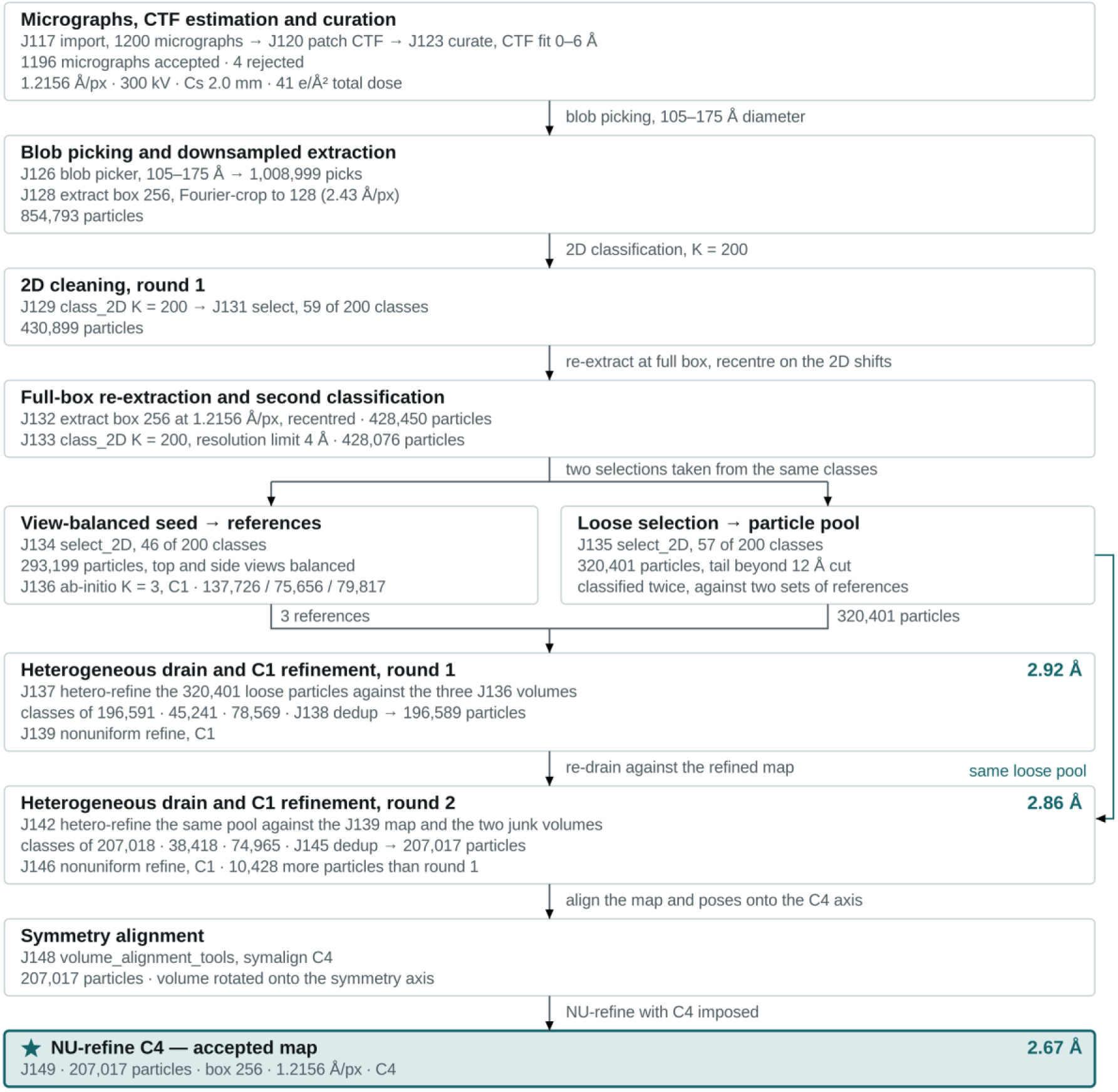
Processing lineage of the TRPV1 reconstruction (EMPIAR-10059).

**Supplementary Figure S5.**
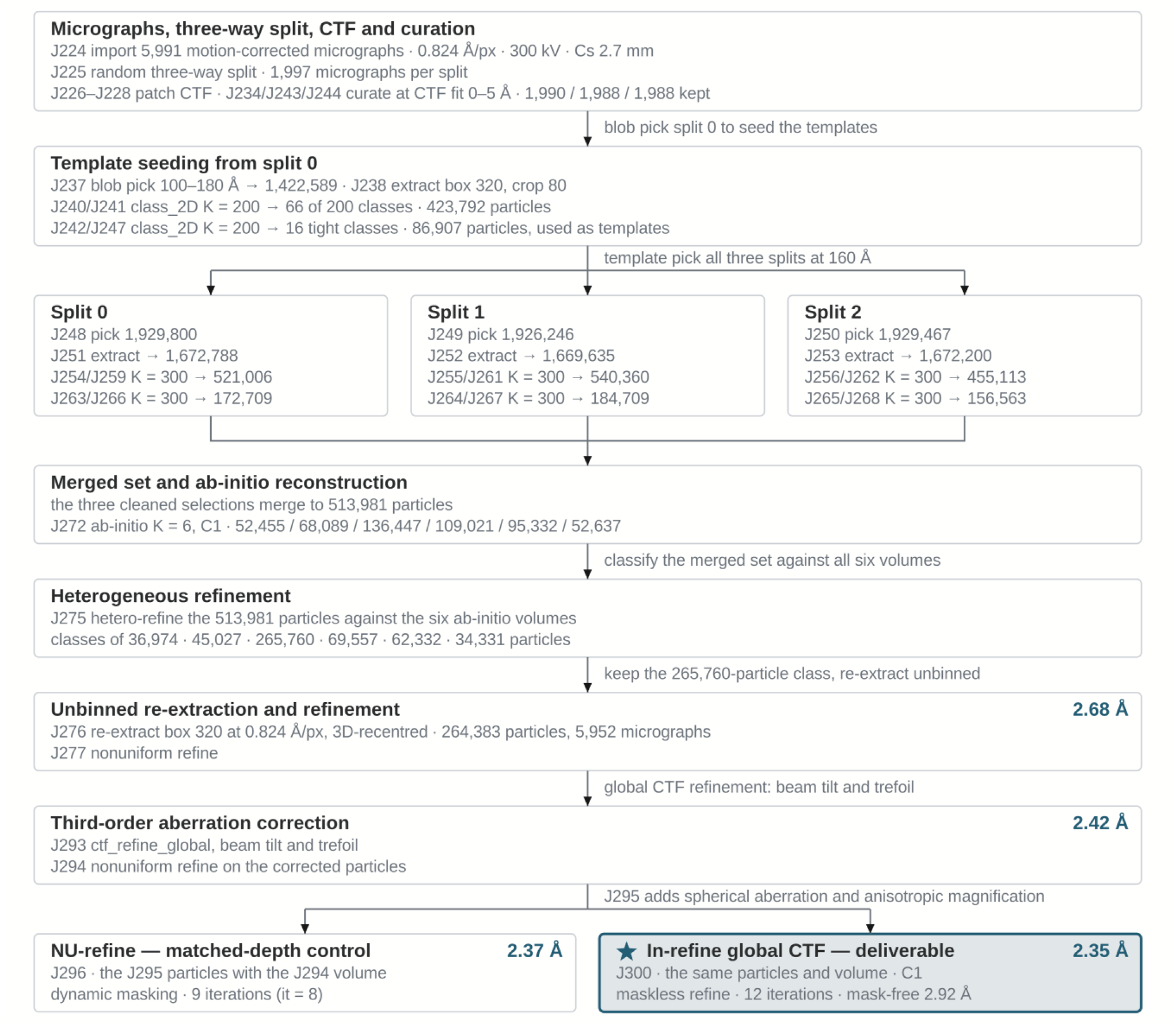
Processing lineage of the GCGR–Gs complex reconstruction.

**Supplementary Figure S6.**
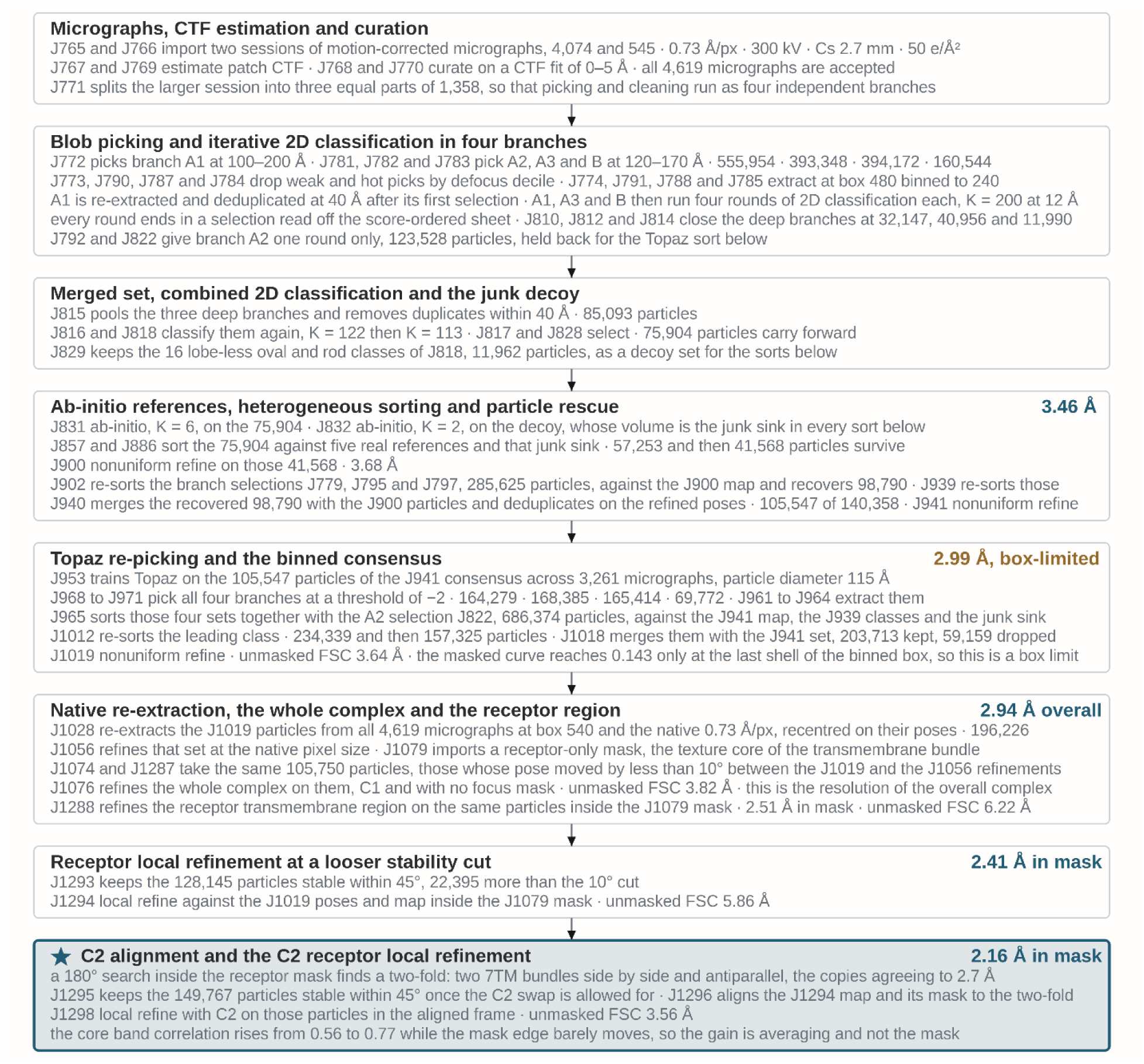
Processing lineage of the somatostatin receptor 5 (SSTR5) reconstruction.

**Supplementary Figure S7.**
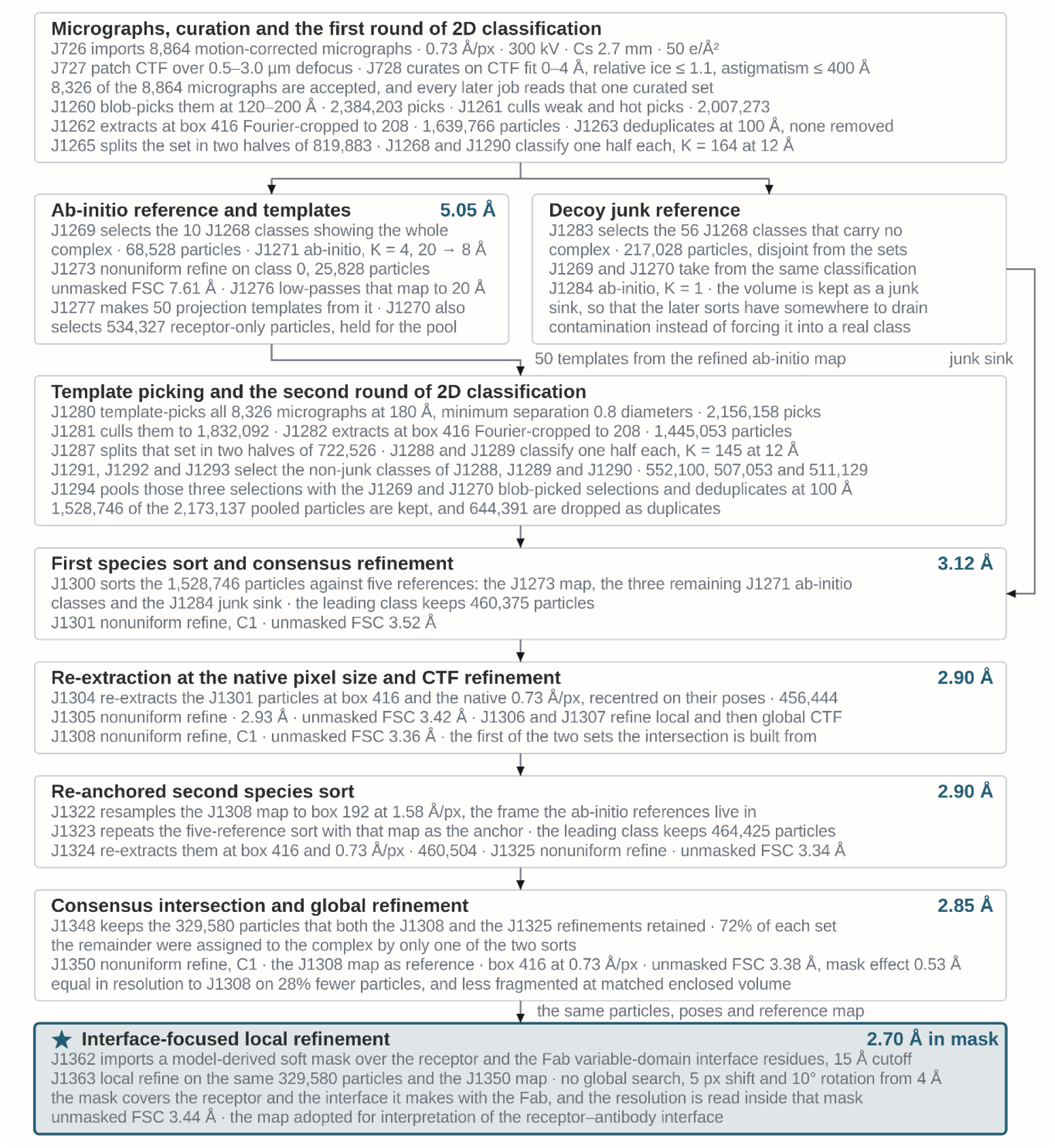
Processing lineage of the cannabinoid receptor 1 (CB1) reconstruction.

**Supplementary Figure S8.**
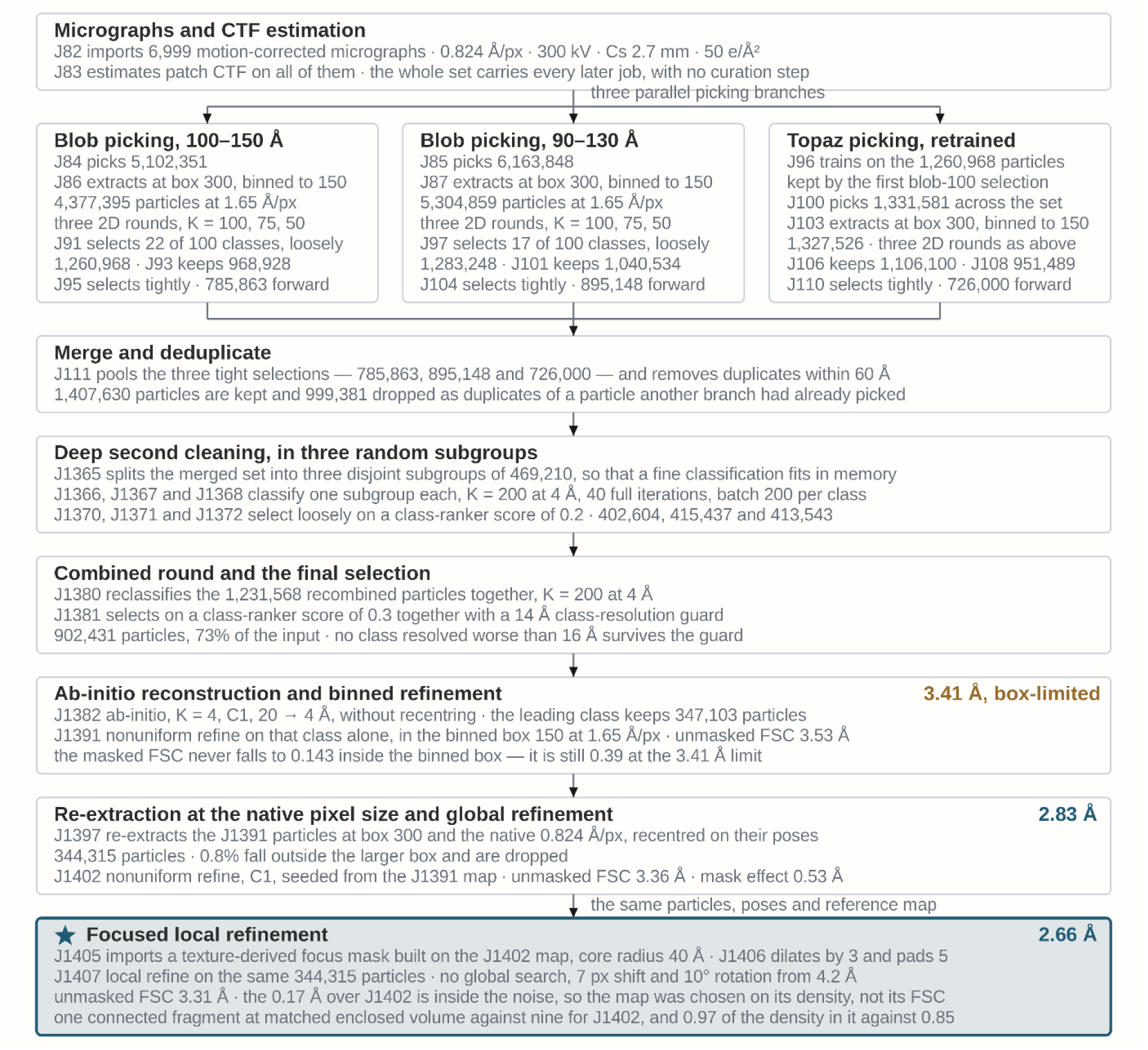
Processing lineage of transporter A (unpublished) reconstruction.

**Supplementary Figure S9.**
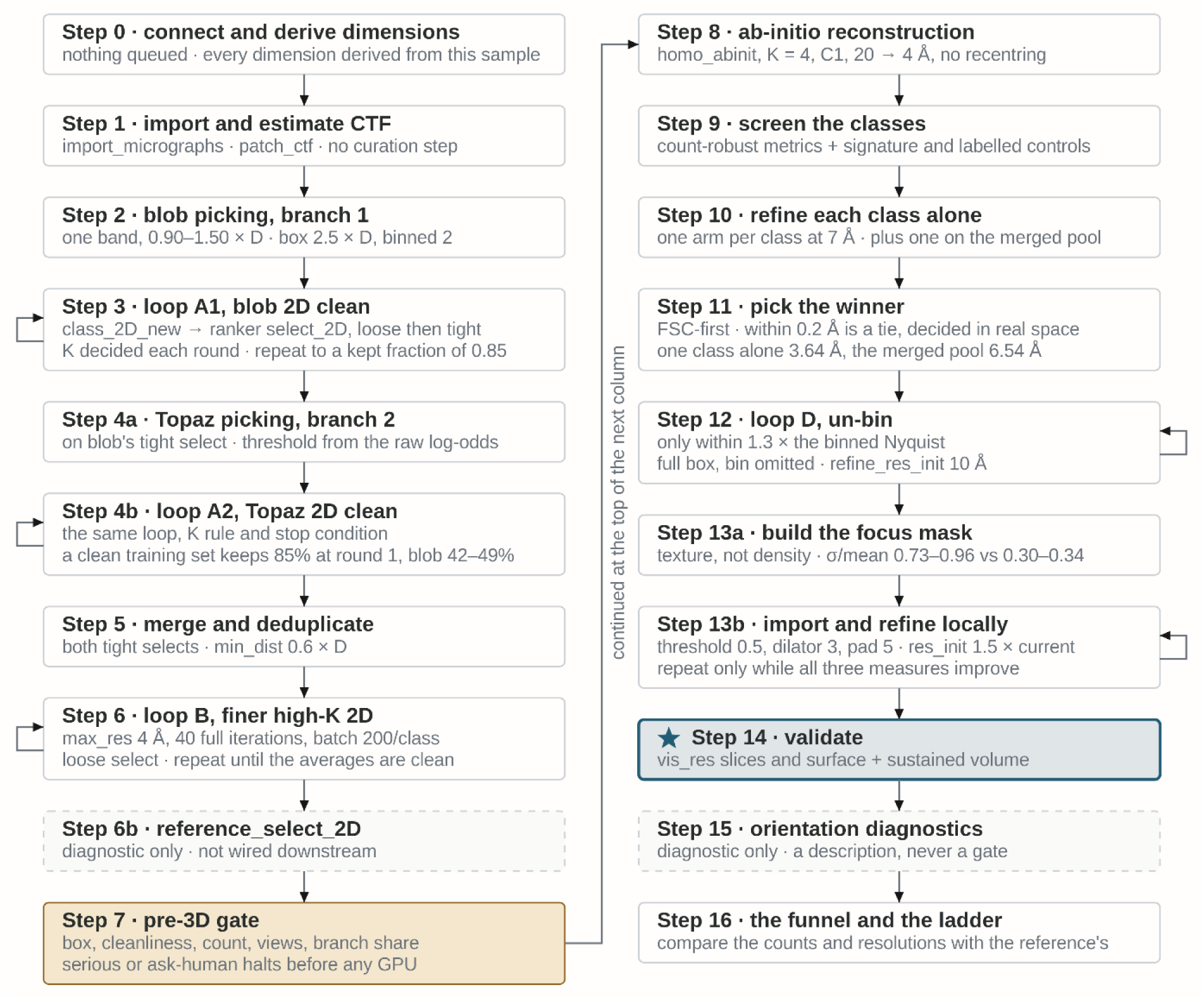
Reusable automatic processing pipeline derived from reconstruction of transporter A, which was used to resolve transporter B (unpublished).

**Supplementary Figure S10.**
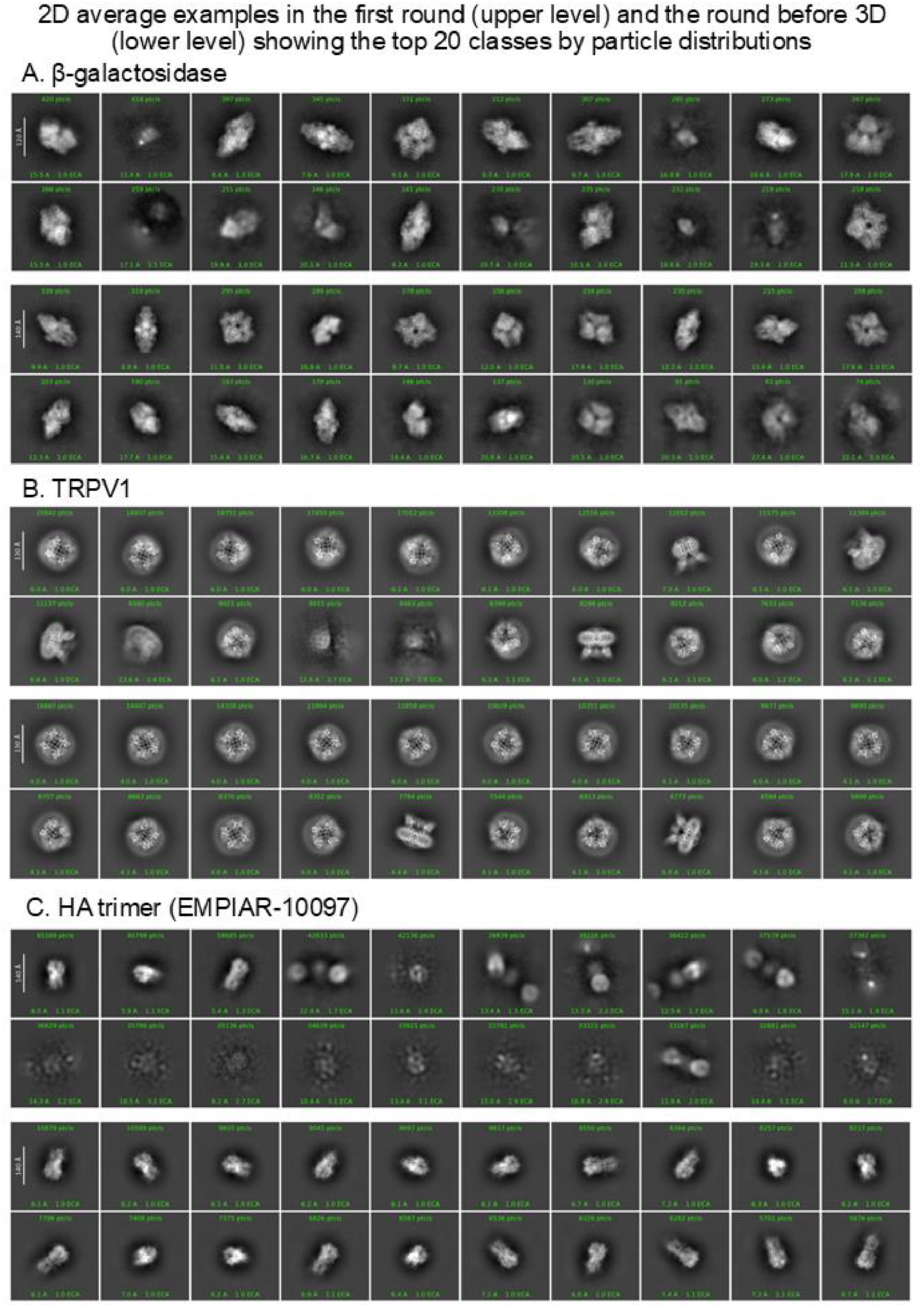
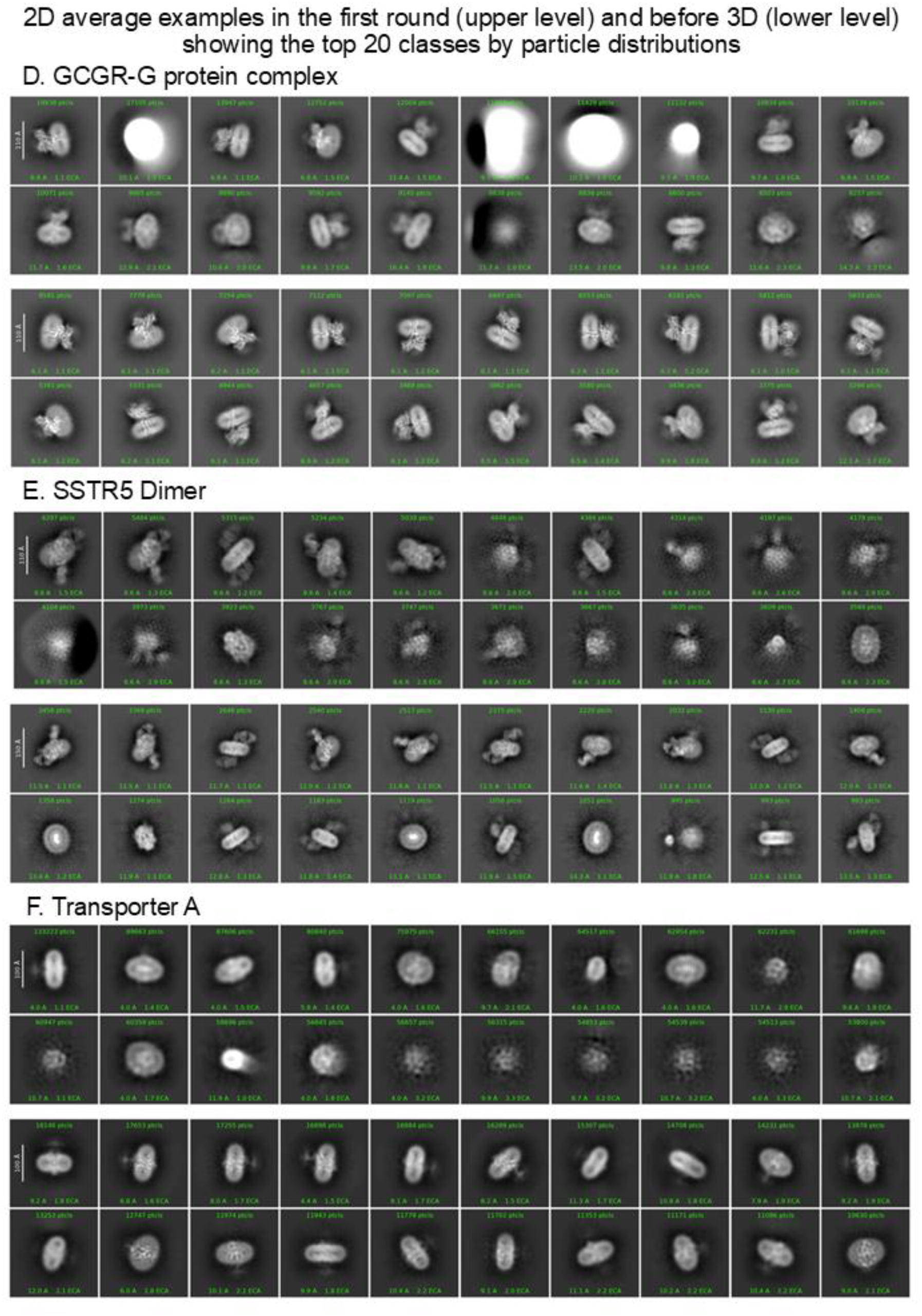
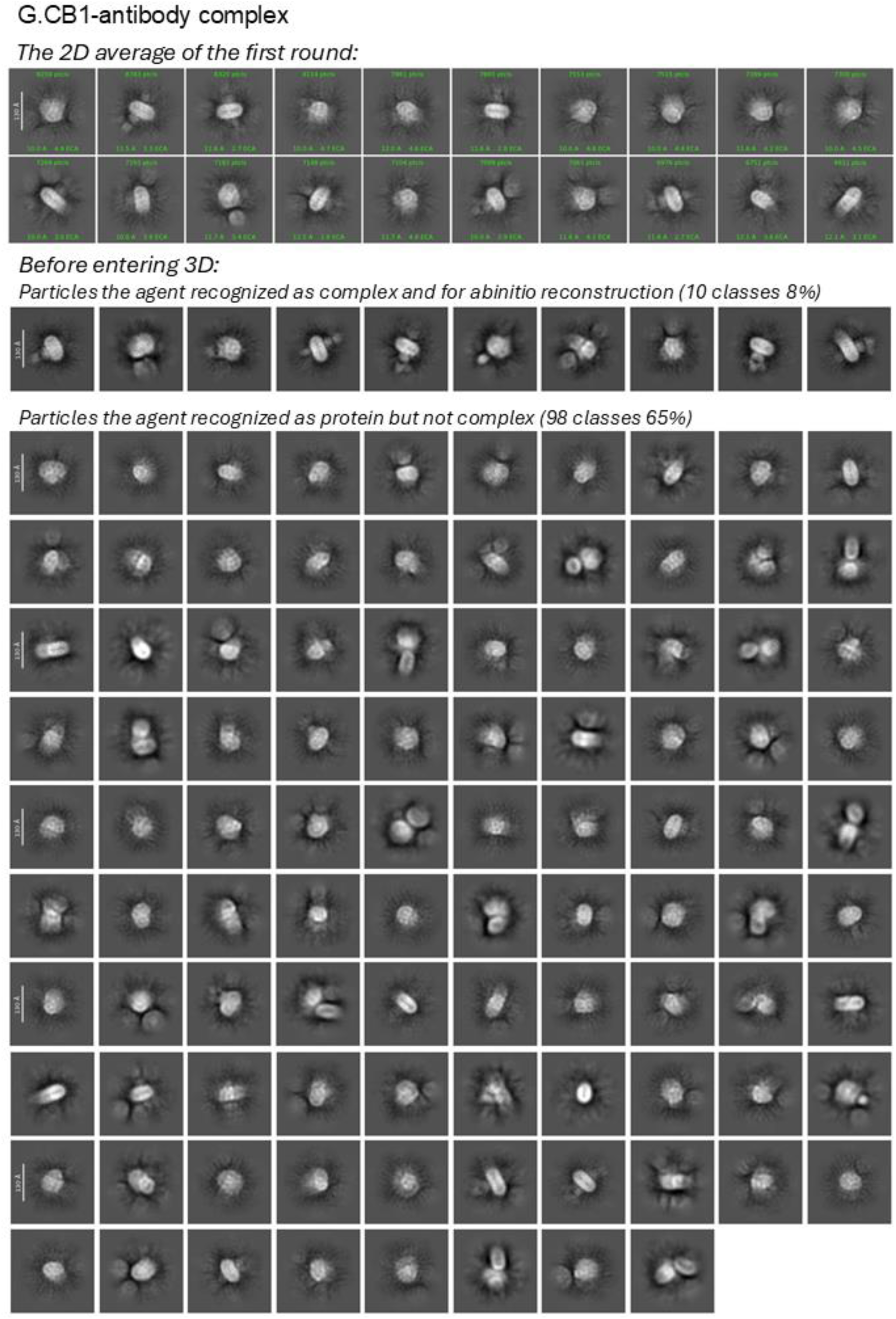
Examples of 2D averages.

**Table S1.** information of the results for tested datasets

| Dataset | Software | Resolution (Å) | Jobs in lineage | Compute time (h) | Human Intervention<sup>*</sup> |
|---|---|---|---|---|---|
| **β-galactosidase, instance 1** | CryoSPARC | 3.02 | 24 | 1.9 | No |
| **β-galactosidase, instance 2** | CryoSPARC | 2.83 | 56 | 2.7 | No |
| **β-galactosidase, instance 3** | CryoSPARC<br>+ RELION | 2.86 | 28 | 1.9 | Yes |
| **Haemagglutinin trimer, tilted (EMPIAR-10097)** | CryoSPARC | 2.86 | 53 | 43.5 | No |
| **Haemagglutinin trimer, untilted (EMPIAR-10096)** | CryoSPARC | 2.88 | 40 | 4.9 | No |
| **TRPV1 (EMPIAR-10059)** | CryoSPARC | 2.67 | 20 | 7.7 | No |
| **GCGR–Gs** | CryoSPARC | 2.35 | 40 | 11.6 | No |
| **SSTR5** | CryoSPARC | 2.16 | 82 | 31.9 | Yes |
| **CB1–Fab** | CryoSPARC | 2.70 | 43 | 32.0 | Yes |
| **Transporter A** | CryoSPARC | 2.66 | 44 | 33.3 | Yes |
| **Transporter B** | CryoSPARC | 2.81 | 45 | 36.8 | No |

<sup>\*</sup>Human interventions related to the reconstruction results:
β-galactosidase (instance 3):
1.Convert the output of J223 to relion and run class3D. (for testing pyem)
SSTR5:
1.Apply C2 on local refinement.
CB1:
1.J1273 is our target protein, go ahead for enrich the particles and push the resolution.
2.your templated picked particle without any 2d class and selection, you intent to it or just missing?
3.also try the merge of J1325 and J1308.
4.the interaction between receptor and antibody is what we concern.
5.for mask making, I suggest based on built model.
Transporter A:
1.2D with finer parameters as J112. (After the retrospecting the project, the operator thinks this suggestion was with some help, but it is not the key for the success)

